# Epigallocatechin-3-gallate inhibit the protein arginine methyltransferase 5 and Enhancer of Zeste homolog 2 in breast cancer both *in vitro* and *in vivo*

**DOI:** 10.1101/2024.02.18.580855

**Authors:** Kirankumar Nalla, Biji Chatterjee, Jagadeesh Poyya, Aishwarya Swain, Krishna Ghosh, Archana Pan, Chandrashekhar G Joshi, Bramanandam Manavathi, Santosh R Kanade

## Abstract

Histone methyltransferases are selectively catalyzing the methylation of lysine or arginine residues of target histone and non-histone proteins, classified as lysine methyltransferases and arginine methyltransferases. The EZH2 and PRMT5 catalyze trimethylation of H3 at K27 and symmetric dimethylation of H4 at R3 respectively. These histone repressive marks have been considered as hallmarks in cancer. Both PRMT5 and EZH2 over expressed in several cancers and have been considered as important target of drug development. As a result, many synthetic molecules as inhibitors of both PRMT5 and EZH2 are at different level of preclinical and clinical phases. Cancer atlas data analysis revealed that both PRMT5 and EZH2 had shown more than 90% amplification in breast cancer alone. We screened an array of phytocompounds towards the inhibition of PRMT5 and EZH2 using *in silico, in vitro* assays. Among them Epigallocatechin-3-gallate (EGCG) has interacted with human PRMT5: MEP50 and EZH2 efficiently. The EGCG interacted within the SAM binding site, with a π-cation interaction at Lys 333 and H-bonds with Tyr324, Tyr334, Gly365, Leu437, and Glu444. Surface plasmon resonance analysis revealed that EGCG has strong binding affinity in nanomolar concentrations with both PRMT5-MEP50 than EZH2. Further in vitro methylation and cell-based assays proved the inhibitory potential of EGCG by reducing the catalytic products of PRMT5 and EZH2 i.e., H4R3me2s & H3K27me3 respectively and showed that it induced autophagy and apoptosis. Furthermore *in vivo*, mouse xenografts studies demonstrated that oral dosage, reduced tumor size significantly with reduction in proliferation marker Ki67 and these histone repressive marks. Finally conclude that inhibition of PRMT5 and EZH2 by EGCG potentially can be used to develop combined therapeutic approaches.

## Introduction

According to Globocan’s recent data, breast cancer has exceeded lung cancer as one of the most diagnosed cancers, with an estimated 2.3 million fresh cases, followed by lung cancer (11.4%), colorectal cancer (10.0%), prostate cancer (7.3%), and abdomen cancer (5.6%). The estimated 2.3 million breast cancer cases represent one out of every eight cancers diagnosed in 2020 **^(1–2)^**. The factors responsible for the occurrence and development of breast cancer include obesity, age, physical activity, family history, and overuse of oral contraceptives. Breast cancer progression involves the accumulation of aberrant changes at both genetic and epigenetic levels, which eventually lead to tumorigenesis. **^(3)^** The epigenetic regulation like DNA methylation, and histone post-translational modifications like methylation, acetylation, phosphorylation, ubiquitination, and sumoylation, alters the architecture of chromatin. These changes are well known to modulate various molecular, cellular, and biological pathways associated with breast cancer progression.

Histone methyltransferases (HMTs) are selectively catalyze the methylation of lysine or arginine residues of target histone or non-histone proteins, and classified as lysine methyltransferases (KMTs) and arginine methyltransferases (PRMTs) respectively. The KMTs are in two families: containing SET domain KMTs, which include Trithorax, Enhancer of Zeste (EZH), Su(var)3-9, and non-SET domain-containing KMTs, like DOT1-like proteins **^(4)^**. The EZH1, and EZH2 are two well-known methyltransferases among KMTs. The PRMTs, include type I, II and III **^(5)^**. Type I PRMTs forms the asymmetric dimethylation of arginine residues that include PRMT1,2,3,4,6,8 whereas type II PRMT5/7 adds the methyl groups to arginine side chain symmetrically while type III only produces monomethylated arginine **^(7)^**. PRMT5, always associated with its activity partner methylosome protein 50 (MEP50), is the predominant member of the type II PRMTs. Both PRMT5 and EZH2 extensively gained attention as a novel drug target and recognized as an oncogene. The EZH2 is catalytic subunit of polycomb repressive complex (PRC2) that leads to the trimethylation of Lys-27 in histone 3 (H3K27me3) which can alter the target genes expression. EZH2 also methylate non-histone protein substrates like transcription factor like GATA4, c-Myc and directly interact with other non histone targets like with NF-κB component, TCF, β-catenin, and cyclin D1 to activate downstream targets. It has critical role in autophagy, apoptosis **^(6)^** cell cycle progression, cellular senescence **^(38)^** DNA damage repair and related signaling pathways. The H3K27me3 signature is regarded as very important epigenetic event during cell fate determination and development. The relations between EZH2 and cancer initiation, progression, metastasis, drug resistance, metabolism and immunity regulation are extensively documented. The abnormal expression of EZH2 gene has been noticed in diverse types of cancer with poor clinical outcomes in breast cancer, prostate, gastric cancer, esophageal cancer, anaplastic thyroid carcinoma, nasopharyngeal carcinoma, and endometrial carcinoma hence EZH2 has been considered as an important target for cancer therapy**^(38)^**. The PRMT5-MEP50 complex consists of four PRMT5 and MEP50 molecules this association very critical for optimal activity **^(10–11)^**. The role of PRMT5 has been well established in tumorigenesis, RNA metabolism, transcriptional regulation, DNA repair, signal transduction and ribosome biogenesis **^(5,10)^**. According to various studies reported, PRMT5 is involved in a multitude of cellular processes, exerting its influence through the methylation of a diverse array of both cytoplasmic and nuclear substrates. Many non-histone proteins like E2F1, KLF4, TP53, EGFR, BCL6, FOXP3, SnRNP assembly (SmB/B’, SmD1, SmD3), CRAF, RAD9 are reported to be substrates of PRMT5 **^(10,14)^**. PRMT5 has been found to methylate specific residues, of histone H4 and H3 residue, H3R8me2s and H4R3me2s are generally together with transcriptional repression in contrast with H3R2me2s associated with transcriptional activation of target genes **^(10,14)^**. The PRMT5-mediated methylation of protein/s is implicated in the control of normal and pathological status, in maintenance of tissue homeostasis, development, survival and self-renewing capacity of stem/progenitor cells in hematopoietic, nervous, muscular, and reproductive systems. Apart from that PRMT5 is upregulated in several human cancers, including breast cancer, lung cancer, prostate cancer, lymphomas and colorectal cancer **^(5,6,10)^**, its overexpression observed in autoimmune diseases **^(12)^**, inflammation diseases **^(13)^** as well. The mechanism of upregulation of PRMT5 in disease status is still unknown. Interestingly recently studies reported that the functional association of both PRMT5 and EZH2 and it has been targeted for combined therapy **^(4,8)^**. Considering fact that PRMT5 and EZH2 are involved in a wide range of physiological processes, and are crucial for the maintenance of cancer phenotypes, and perturbation of PRMT5 and EZH2 may be a novel strategy. The methyltransferases being dependent on the cofactor (methyl group donor) SAM or AdoMet were targeted with SAM-analogue compounds or structural derivatives, mostly of synthetic origin **^(40)^**. Many researchers are targeting to find the small molecular inhibitors to develop inhibitors against PRMT5 **^(4,10,14)^**. EZH2 **^(4, 8,9)^**. A large number of selective SAM-competitive inhibitors of EZH2 have been developed to inhibit both the wild and mutant form of EZH2 **^(39,41)^**. The SAM-competitive molecules, like CPI-0209, Tazemetostat (EPZ-6438), CPI-1205, PF-06821497, and DS-3201, a dual inhibitor of EZH1 and EZH2, have entered into clinical trials **^(^**^37, 38^**^)^**and there are several reported inhibitors for EZH2, but very few like three only completed phase-1 and 2 trails successfully, Tazemetostat (EPZ6438)-NCT0310982, Atezolizumab-(NCT02220842), NCT02875548 **^(15)^** which are under phase-1 clinical trials **^(10,14,35,36)^**. Like EZH2, PRMT5 also has been targeted by small molecular inhibition. The synthetic SAM analogue, sinefungin, GSK-3326595, EPZ015666, showed very good inhibition of PRMT5 by targeting to SAM or peptide binding site of PRMT5 **^(42)^**. The GSK3326595 an improved form of EPZ015938, entered in clinical trial alone or in combination pembrolizumab (NCT02783300) or 5-azacytidine. (GSK-3326595[NCT02783300], JNJ-64619178[NCT0357310] and PF06939999 [NCT03854227]**^(10,14)^**. Given the significant importance of PRMT5 and EZH2 as critical targets for anticancer therapies, the limited pool of known inhibitors emphasizes the necessity for increased attention and effort toward the development of inhibitors specifically targeting PRMT5 and EZH2. Natural molecules have an advantage over chemical compounds because of their long-term use without or minimum side effects.

The EGCG, is the major and most abundant catechin present in green tea, chemically, belongs to the big family of plant-derived polyphenol with a strong antioxidant activity **^(16)^**. EGCG have been shown to inhibit carcinogenesis by affecting large number of signal transduction pathways and transcription factors involved in premalignant and malignant cells by inducing autophagy, apoptosis cell and cycle arrest, selectively. It also been reported that EGCG inhibit the angiogenesis, induce epigenetic modifications, carcinogenesis, and metastasis in animal models **^(17,18)^**. Plenty of reports in animals, as well as case-control and epidemiologic studies in humans, reveal that green tea consumption or oral administration for long-term significantly reduces the incidence of various cancers **^(19,20)^**, indicated a vital role of EGCG in the chemoprevention of cancer. The large number reports suggested the excellent anticancer potency of EGCG alone as well as along other compounds or drugs, however the specific binding target for EGCG is not been studied extensively. In the present study first time provide ample evidence on binding of EGCG to potential target of cancer PRMT5 with high binding potential using molecular docking, surface plasma resonance (SPR) and its inhibitory potential has been validated in both *in vitro* and *in vivo* studies.

## 2. Materials and Methods

### 2.1 Materials

Epigallocatechin-3-gallate (EGCG), Dimethyl sulfoxide (DMSO), S-adenosyl methionine (SAM), and 3-(4,5-Dimethylthiazol-2-yl)-2,5Diphenyltetrazolium-Bromide (MTT), Recombinant proteins (PRMT5:MEP50 complex (SRP0145-25470) and EZH2 (SRP0379) were purchased from Sigma-Aldrich USA. CM5 Sensor chip for SPR (cat no: 50-105-5511) was purchased from GE Healthcare Bio-Sciences (Cytiva life sciences-USA). The antibodies Anti-PRMT5 antibody (#2252) Histone (H4) antibody (#2935), H3 antibody (#4499), EZH2 antibody (#5246), H3K27 (#9733), anti-beta actin antibody (#4967), Ki67 antibody (#D3B5) was obtained from Cell Signaling Technology (CST) USA. H4R3me2s Polyclonal antibody (A3718-050) from Epigentek NY-USA, FITC-Annexin V Dead cell/ Apoptosis kit (V13242) from Invitrogen-USA. PRMT5 monoclonal antibody (MA-125470), EZH2 monoclonal antibody (MA5-15101) procured from ThermoFisher. The antibodies for PARP (A8770), Cytochrome-c (A0225), BCL-2 (A0208), Bax (A0207), Bad (A1593), Beclin-1 (A7353), LC3 I/II (A19665), ATG5 (A0203), ATG12 (A19610), ATG7 (A19604), ATG3 (A19594), Cyclin-A2 (A7632), Cyclin-B1 (A19037), Cyclin-E1 (A14225), Cyclin-D2 (A1773) procurred from Abclonal USA. All other chemicals were of analytical or higher standards unless otherwise mentioned.

### 2.2. Cancer datasets analysis for role of PRMT5 & EZH2

To find out the role of PMRT5 & EZH2 in cancer, cBioportal dataset **^(61)^** which is an exploratory analysis tool meant for exploring large scale cancer genomic datasets, made up of curated data from the large consortiums like The Cancer Genome Atlas (TCGA) and Therapeutically Applicable Research to Generate Effective Treatments (TARGET), as well as from individual lab publications. In cBioportal site we selected all cancer datasets and proceed for searching by the gene name (EZH2 & PRMT5) and considered parameters like Amplification, mutations, deletions, and multiple alterations as main parameters and retrieved data, analysed, and documented. To explore the correlation between expression of PRMT5 & EZH2 and survival of patients in 30,000 (thirty thousand) samples from 21 tumor types including breast, ovarian, gastric, lung, colon, Acute myeloid leukemia, and myeloma. The cBioportal **^(61)^** dataset the tool which is developed for discovery and validation of survival biomarkers was used. We selected breast cancer patients’ data for plotting Kaplan-Meier curve for both the oncoproteins (PRMT5 & EZH2).

### 2.3 Molecular Docking

The crystal structures of *Homo sapiens* PRMT5: MEP50 complex (with sinefungin (SAM analog) and EZH2 (enhancer of zeste homolog2) determined by X-ray diffraction at 2.35 Å resolution was retrieved from the RCSB Protein Data Bank (PDB ID of PRMT5: 4X60, 4X61, 4X63, EZH2’s PDB-ID: 5HYN & 4Mi5) respectively. Both the proteins were prepared by adding the missing residues using the Prime wizard of Schrodinger. The completed protein structures of PRMT5: MEP50 complex and EZH2 was submitted to Protein Preparation Wizard in the Maestro tool to assign bond order, hydrogen, and disulfide bonds. All seleno-methionines in the structures were converted to methionine residues, and the protein structures were energy minimized using the OPLS3 field2005 force filed used to minimize the energy 4.

For docking, the receptor grid was created around the active site of the proteins, determined by identifying the interacting residues in the reference molecule bound to the crystal structure. Subsequently, all the prepared ligands were docked using Glide (grid-based ligand docking with energetics) program (Schrödinger Release 2019-3: Glide, Schrödinger, LLC, New York, NY, 2019 **^(21)^**, with existing limits (e.g., Van der Waals radii of 1.0) with XP (extra precision). The workflow process involved high throughput virtual screening (HTVS) followed by SP (standard precision), and ultimately, XP mode leading to accuracy, best scoring, and facilitate visualization of the docked ligands. The Glide results were then examined for individual ligand poses to visualize atoms by proximity (5 Å), hydrogen bonds, and other contacts as well as the Glide XP scoring functions.

### 2.4 Surface Plasmon Resonance (SPR) binding studies

Surface plasmon analysis was performed using the Biacore-T-200 system (GE Healthcare). Both recombinant PRMT5-MEP50 complex and EZH2 (50µg) were diluted in 50µL in 1x PBS of pH-7.4 to achieve a final protein concentration of 1mg/mL) / (exposure 30 sec/2µl per minute), and 20µg of protein was immobilized covalently to the hydrophilic carboxymethylated dextran matrix of a CM5 sensor chip by the standard primary amine coupling reaction as described by the manufacturer. The amount of immobilized protein was estimated by assuming that 2500 RU correspond to 2ng of immobilized protein/mm^-2^ and a surface concentration of 2500 RU per 2ng/mm^-2^ was used in the analysis.

The chip surface comprising flow cells (FC) 1 (EZH2) and (FC) 3 (PRMT5-MEP50 complex) served as the reference surface (without protein) and was directly deactivated by injecting 1M ethanolamine (pH 8.5) for 7 minutes. Flow cells 2 and 4 (FC-2 for EZH2, FC-4 for PRMT5-MEP50 complex) were activated by exposure to an immobilization buffer (a mixture of 200 mM N-ethyl-N’-dimethylaminopropyl carbodiimide (EDC) and 50 mM N-hydroxysuccinimide (NHS)) for 7 minutes. A stable baseline was achieved in the cell with immobilized protein by continuously flowing HBS-P running buffer (10 mM HEPES pH 7.4, 150 mM NaCl, 0.005% P20+DMSO buffer filtered through a 0.22 µm filter) at a rate of 5-10 µL/min for 7 minutes. This equilibration step also eliminated any low-affinity ligands bound to the protein as isolated. All measurements were typically conducted at 25°C using running buffer (10 mM HEPES pH 7.4, 150 mM NaCl, 0.005% P20+DMSO buffer filtered through a 0.22 µm filter) at a constant flow rate of 30 µL/min. EGCG was dissolved in the running buffer and analyzed (in triplicate) using a dilution series ranging from two to five folds.

The EGCG was prepared in DMSO as 1M stock and diluted in 1x-PBS (pH7.4) to make 10 mM working stocks and injected over PRMT5-MEP50 complex & EZH2 at 30µl/min for a contact time of 60 seconds at 25°C, three-cycle kinetic analysis was performed for interaction analysis between PRMT5-MEP50 complex, EZH2 with EGCG. After subtracting the buffer blank average, the corrected sensograms were analysed by employing BIA-T200 evaluation software version 2.1 (GE Healthcare). The relation between protein concentration and the Response unit (RU) value was calculated based on the formula, 1000 RU = 1ng/mm^2^ – for surface concentration, 1000 RU = 10mg/ml – volume concentration. The dissociation rate constant (k_d_) was calculated based on Langmuir adsorption model.

### 2.5 Cell culture and cell viability assays

The human malignant cell lines: Du-145 (prostate), A549 (lung), HeLa (cervix), Hep G2 (liver), MCF7 (breast), MDA-MB-231 (breast) PC-3 (prostate) and HEK293 (human embryonic kidney) were acquired from the National Centre for Cell Science (NCCS) cell repository located in Pune, Maharashtra, India. All cell lines were cultured and maintained under standard sterile conditions at low passage numbers. The cells were cultured in sterile cell culture dishes or flasks using a complete medium consisting of high-glucose Dulbecco’s modified Eagle’s medium (DMEM) supplemented with 10% (v/v) fetal bovine serum and 1x antibiotic-antimycotic solution. Culturing was carried out at 37°C in a moistened incubator with an atmosphere containing 95% air and 5% CO2.

#### 2.5.1 MTT assay

The viability assay was conducted briefly: Initially, 5000 cells were seeded per well in triplicates on 96-well plates and allowed to grow overnight. Subsequently, the media were replaced with DMEM alone. After 6 hours of incubation, various concentrations of EGCG were added to the wells, and the cells were further incubated for 24 and 48 hours. Following the respective incubation periods, the MTT assay was carried out according to previously established protocols**^(22)^**. Briefly, MTT solution was added to the wells and allowed to incubate for 3 hours. After aspirating the MTT solution, 100 µL of DMSO was added to each well, followed by colorimetric quantification using a plate reader (Tecan). By analyzing the optical density readings of the test samples and blanks, the percentage viability of the cells was calculated.

#### 2.5.2 Assessment of Cell Viability Using Trypan Blue Dye Exclusion test

Trypan blue dye exclusion test was according to previously established protocol **^(22)^**. The experiment involved combining a cell suspension post-EGCG treatment with dye, followed by microscopic examination to determine whether cells absorbed the dye or excluded it. A graph was constructed based on the percentage of viable cells, characterized by clear white cytoplasm, versus non-viable cells, identified by blue cytoplasm. The percentage of cell viability was calculated using the formula:

% Cell Viability = Abs (Test sample) / Abs (Control) × 100.

The percentage of cell inhibition was then determined as:

% Cell Inhibition = 100 - Percentage of Cell Viability.

#### 2.5.3 Cellular Morphology with EB/AO Double Staining

The Ethidium Bromide and Acridine Orange double staining assay was conducted following established procedures as described previously **^(22)^**. This method employs Acridine Orange as a viability stain, which permeates into all cells, and Ethidium Bromide, a non-permeant dye that penetrates only compromised cell membranes, indicating cell necrosis following apoptosis. While cells with Acridine Orange emit green fluorescence, dead cells that have absorbed Ethidium Bromide display red fluorescence, with necrotic cells appearing orange and resembling viable cells, albeit with minimal chromatin changes. After incubating MDA-MB231 cells with EGCG for 24 hours, they were labeled with EB/AO and observed under a fluorescent microscope.

#### 2.5.4 Colony formation assay

The colony formation assay was performed by according to **^(23)^**. with little modifications. MCF-7 cells (500 cells per dish) seeded in 30mm dishes and allowed cells to attach to the plates, once cells were attached to plate, cells were given treatment with EGCG with following concentrations (0, 0.1, 1 & 10 µM) and incubated for 24 hours, after incubation, replaced with serum free media and allowed to grow them until cells in dishes have formed sufficient colonies. Once colonies were appeared in dishes, removed media given PBS wash carefully, and added 2-3 ml of 6% glutaraldehyde and 0.5% crystal violet, incubated for 30 minutes, removed crystal violet & glutaraldehyde mixture, and rinsed with tape water by filling the dishes, kept for air dry at room temperature for 20 minutes and triplicate culture dishes were maintained for the experiment and counted the colonies and plotted the graph as the percentage of colony number.

#### 2.5.5 Apoptosis and Cell cycle studies

Cells were initially seeded in six-well plates at a density of 1.5 million cells per well, followed by treatment with EGCG at concentrations ranging from (0.1 to 10 µM). The cells were then incubated for 24 hours. After the incubation period, the cells were trypsinized and stained with Annexin V-fluorescein isothiocyanate (FITC) at a dilution of 100x and propidium iodide (PI) at a concentration of 0.5 µg/ml for 15 minutes at room temperature. Subsequently, the fluorescence intensities of Annexin V-FITC and PI were analyzed using FACScan (Becton Dickinson, San Jose, CA, USA). Gating was performed as follows: Annexin V (+)/PI (-) cells were identified as apoptotic cells, Annexin V (+)/PI (+) cells as undergoing secondary necrosis, and Annexin V (-)/PI (+) cells as necrotic cells. The obtained results were analyzed using FlowJo software.

For cell cycle analysis following treatment with the aforementioned concentrations, cells were trypsinized and fixed in 70% ethanol overnight at -20°C. They were then stained with RNase-A at a concentration of 100 µg/ml and Propidium Iodide (PI) at a concentration of 50 µg/ml for 15 minutes and subjected to flow cytometric analysis using FACScan (Becton Dickinson; San Jose, CA).

#### 2.5.6 Acidic Vesicular Organelle (AVO) staining by Acridine orange

Acridine orange (AO) permeates cell and organelle membranes freely, serving as a crucial indicator for acidic vesicular organelle (AVO) formation. These AVOs emit green fluorescence throughout the cell, with the exception of acidic compartments, primarily late autophagosomes, where they emit red fluorescence. This distinctive characteristic of AVO development is particularly evident during autophagy, signaling autophagosome maturation and an efficient autophagic process, as only mature/late autophagosomes exhibit acidity. The intensity of red fluorescence correlates with the abundance of AVOs in autophagic cells. To assess this, cells were treated with specified concentrations of EGCG, followed by removal of the media and addition of fresh media containing 5 ug/mL acridine orange. Subsequently, cells were incubated for 10 minutes at 37°C and observed under a fluorescent microscope.

### 2.6 Western blotting

The cells underwent treatment with different concentrations of EGCG (0, 0.1, 1, and 10µM) for the specified duration. Subsequently, they were rinsed with phosphate-buffered saline and lysed using RIPA buffer containing a protease inhibitor cocktail, 1 mM DTT, and 1 mM PMSF at 4°C. The resulting cell lysate underwent centrifugation at 10,000 rpm for 10 minutes at 4°C. Protein concentration was determined using Bradford’s protocol with BSA as a standard, and equal amounts of proteins were loaded onto a 10% SDS-PAGE gel following Laemmli’s method. The proteins were then transferred onto nitrocellulose membranes for immunoblotting. The membrane was blocked with skimmed milk in Tris-buffered saline containing tween-20, and primary antibodies against the target proteins were applied at varying dilutions, followed by overnight incubation. A horseradish peroxidase (HRP)-coupled anti-rabbit secondary antibody at a 1:10000 dilution was utilized. Western blot developing solutions from Bio-Rad were employed, and band intensity was assessed using a Versadoc Imaging System with Image Lab 5.1 software (Bio-Rad). β-actin was used as a loading control.

#### 2.6.1 Preparation of lysate from tumor samples

After dissection, tumor tissues from both the treated and control groups were weighed and washed with chilled 1x PBS and chopped the tissue into smaller pieces whilst keeping it on ice, added 300μl of ice-cold RIPA buffer with protease inhibitor and homogenized with an electric homogenizer with constant agitation for 1h at 4^0^C, after homogenization, centrifuged for 20 minutes at 12,000 rpm at 4°C, the tubes were carefully removed and placed on ice. The supernatant was then collected in a fresh tube kept on ice, while the pellet was discarded. Protein concentration was determined using the BCA kit method, and the samples were subsequently prepared for western blot analysis following the procedure outlined in section 2.6.

### 2.8 Animal studies

Four-week-old Swiss nude mice were procured from the Centre for Cellular and Molecular Biology (CCMB) laboratories in Hyderabad and housed in the animal facility at the University of Hyderabad. All experimental procedures involving animals were approved in advance by the institutional ethics committee at the University of Hyderabad, in accordance with the university’s guidelines for the care and use of laboratory animals. The animal cages were maintained at a temperature of 22±4°C with humidity ranging from 50% to 60%, under a 12-hour light/dark lighting cycle. The mice were provided with a standard laboratory diet and water ad libitum and were allowed to acclimate for one week. Following the acclimatization period, the mice were randomly divided into three groups, each consisting of seven mice: (1) Control group 1, (2) Control group 2, and (3) Treatment group.

#### 2.8.1 Tumor xenograft

MDA-MB-231 cell lines sourced from ATCC were utilized to establish xenografts following previously established protocols **^(26)^**. Nude mice were subcutaneously injected with MDA-MB-231 cells (2x10^6) on both flanks. Tumor volume was assessed every three days using digital vernier calipers and calculated using the formula: Tumor volume = (width^2 x length)/2. Upon reaching a volume of 100 mm^3, EGCG (100 mg/kg) was administered for 21 days. Tumor samples were then collected for weighing and subsequent analysis in further experiments.

### 2.9 Immunohistochemical assays

Hematoxylin and Eosin (H&E) staining were performed according to established protocols^27^. Tumor tissues from both experimental and control mice were fixed overnight in 10% neutral buffered formalin, followed by embedding in paraffin and sectioning. H&E staining has long been utilized by pathologists for routine diagnosis due to its effectiveness in visualizing cellular and tissue structures. Deparaffinization of mouse tissues involved treatment with xylene and rehydration in a series of ethanol solutions of decreasing concentrations for 2 minutes each, followed by a 1-minute water wash. Subsequently, slides were stained with hematoxylin for 3 minutes, followed by another 1-minute water wash, differentiation with malic acid for 1 minute, and then a thorough wash in tap water to remove residual alkali. Counterstaining with eosin lasted for 45 seconds, followed by washes in ethanol and xylene. The slides were covered with a coverslip and observed under a microscope.

For additional histochemical analysis, mouse tissues were harvested, embedded in paraffin, and sectioned. Deparaffinization of experimental mouse-derived slides followed a similar procedure using xylene and varying concentrations of ethanol solutions. After a wash with PBS, tissue sections were blocked with 10% BSA in PBST for 1 hour at room temperature, followed by overnight incubation with anti-Ki67 (proliferative marker from Abcam, diluted 1:500) at 4°C. After PBS washes, the sections were incubated with an anti-mouse secondary antibody (CST, diluted 1:100) for 2 hours, followed by TBS washes. Tissue sections were then stained with DAB (Vector Laboratories Inc.) and visualized under a microscope (Leica, Germany).

### 2.12 Statistical Analysis

Statistical significance was assessed using analysis of variance (ANOVA), with a significance level set at P < 0.05. This threshold was utilized to determine the significance of both treated and untreated samples.

## 3. Results

### 3.1 Exploring the role of PRMT5 & EZH2 in Cancer from database

*In-silco* analysis of PRMT5 & EZH2 role in cancer by using databases cBio-portal and TCGA was carried where 10851 & 10990 number of patients was reported with 11632 and 10960 number of samples in 25 studies respectively. Analysis revealed that both PRMT5 & EZH2 was reported in many cancers, and among all cancers PRMT5 & EZH2 amplification was more than the mutation and deletion (Fig1a-1b). Further analysis of PRMT5 & EZH2 amplification to find the absolute frequency of amplification in different types of cancers showed that the Breast cancer (96% frequency), Lung adenocarcinoma (35%), Pancreatic adenocarcinoma (29%) and Urothelial bladder carcinoma (28%) for PRMT5, whereas for EZH2 only amplification was more in breast cancer alone compared with Pancreatic adenocarcinoma (48%) Lung adenocarcinoma (32%), and Urothelial bladder carcinoma (12%). (Fig1c-1d). Oncoprint visualization of PRMT5 & EZH2, revealed that both major type of alterations in cancer patients due to amplification of PRMT5 & EZH2 (Fig1e-1f).

**Figure-1:**
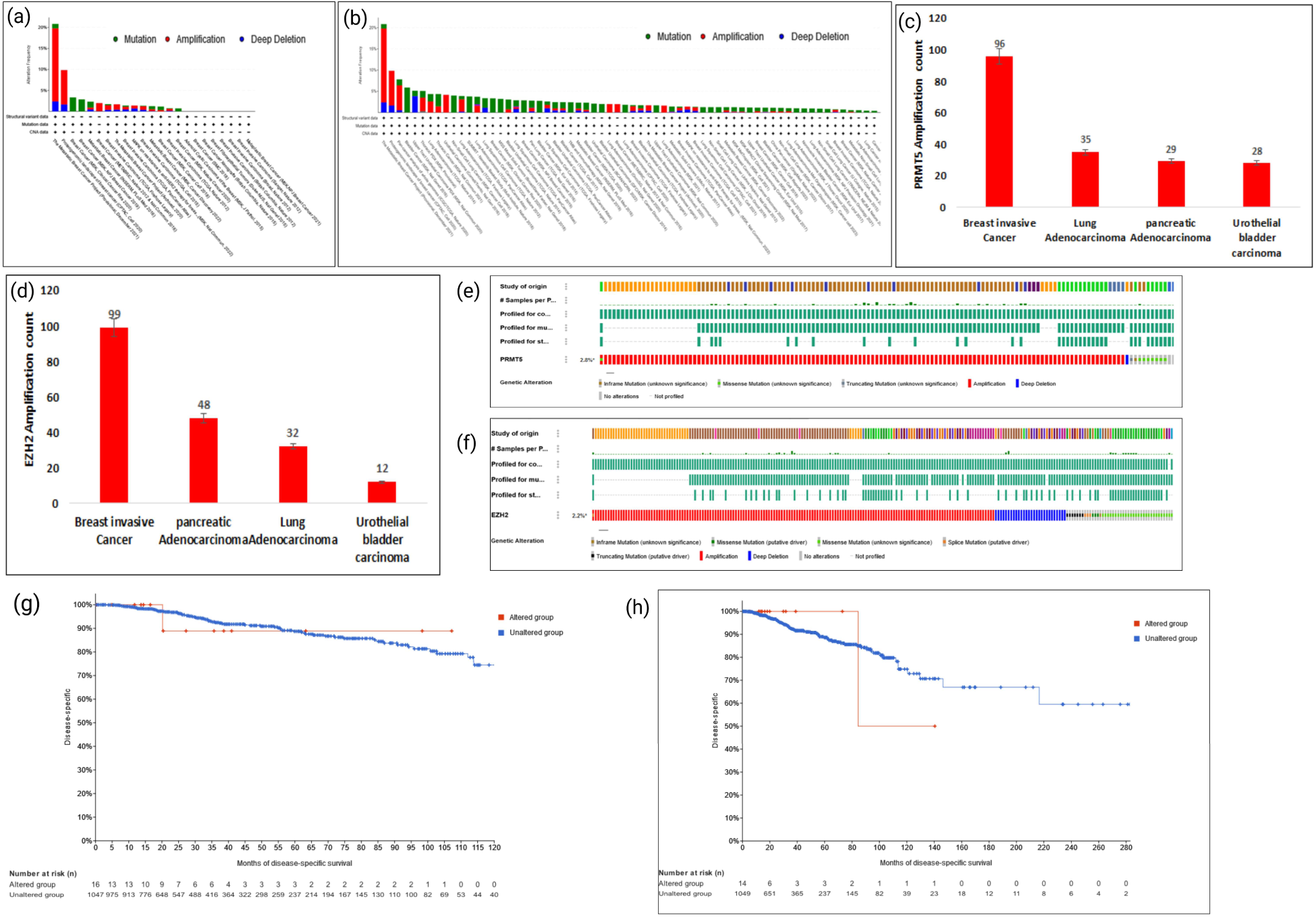
cBioportal data for exploring the role of PRMT5 & EZH2 in cancer database: (a-b) Cancer types summary of PRMT5 and EZH2 in all cancer types, and red color indicates the amplification which is more in breast cancer datasets for PRMT5 & EZH2. (c-d) The amplification frequency data of PRMT5 & EZH2 in breast, lung, pancreatic and urothelial bladder cancers. Breast cancer has highest amplification count for both PRMT5 & EZH2. (e-f) Oncoprints visualization of PRMT5 & EZH2, revealed that both major type of alterations in cancer patients due to amplification of PRMT5 and EZH2. (g-h) Survival analysis of PRMT5 and EZH2 gene by Kaplan Meier curve: we clearly observe that high PRMT5 & EZH2 expression is associated with poor patient survival.

### 3.2 PRMT5 is a negative prognostic factor in breast cancer patients

We constructed Kaplan Meier plot for breast cancer to check the correlation between PRMT5 & EZH2 and patient survival in breast cancer. We used cBioportal dataset and observed that high PRMT5 and EZH2 expression is associated with poor patient survival (Fig1g-1h). It is worth noting that PRMT5 overexpression is particularly in strong association with poor patient survival compared to EZH2. These observations supported previously reported studies of both PRMT5 & EZH2 in alone and in combination, as their role as critical tumor promoter. Analysis reveals and reminds the therapeutic significance of inhibiting PRMT5 & EZH2 in cancers.

### 3.3 Screening of potent PRMT5-EZH2 inhibitors

A library of more than 1200 natural molecules were collected from the literature on different plants which includes majority of them are phenolic or polyphenolic molecules (879), alkaloids, organosulfur compounds, as well as carotenoids (data not shown). The canonical SMILES of these molecules was retrieved from PubChem or ZINC databases and used for ADME analysis in the SwissADME web tool. The filtration criteria to comply with Lipinski and Veber rule for drug-likeness resulted in more than 300 molecules (with the screening of phytocompounds with the binding potential and *in*-silico profiling revealed that a few molecules with high potential bind to catalytic site of PRMT5-MEP50 complex and EZH2. no violations), and more than 20 molecules (with few violations what these violations) were allowed for restricted use in the molecular docking. Each of the ligands was prepared using LigPrep, and the pharmacophore on the PRMT5: MEP50 complex & EZH2, was developed using the Schrödinger workflow Finally, these molecules were docked using the Glide XP program.

### 3.5 Interactions of PRMT5 and EZH2 with EGCG

The ligand-protein docking revealed that EGCG(Fig S1) was interacting with human PRMT5-MEP50 complex & EZH2. The binding pattern of EGCG was analyzed in all the three forms of structures of PRMT5-MEP50 (4X60, 4X61, 4X63) where ligand is SFG, SAM and SAHA respectively (Fig2a-2c) and with EZH2 (5HYN, 4mi5) where ligand is SAHA and SET domain respectively (Fig2d-2e). As shown in the Figure 2, EGCG has similar interactions as SFG, SAM and SAH, 5HYN, & 4mi5. The SFG interacted with PRMT5 protein at the Tyr324, Glu392, Asp419, Met420, Glu435, Glu444 with H-bonds, and a π-cation interaction at Lys393 position. SAHA interacted with EZH2 protein at the Trp624, Ile 109, Ala733, His689, Asn688 and Arg685. Interestingly, EGCG interacted within the same pocket involving a π-cation interaction at Lys 333 and at least five H-bonds with the residues (Tyr324, Tyr334, Gly365, Leu437, and Glu444) in the PRMT5 pharmacophore. Similarly, with 5HYN (EZH2), interacted with five hydrogen bonds (Trp624, Ile109, Ala733, Asn688 and His689), with 4mi5 Leu671, Asn698, Ser669 and Tyr731. The common interactions of EGCG all structural forms of PRMT5 (4X60, 4X61, 4X63) and EZH2 (5HYN, 4mi5) and docking score were tabulated in (Table 1).

**Figure-2:**
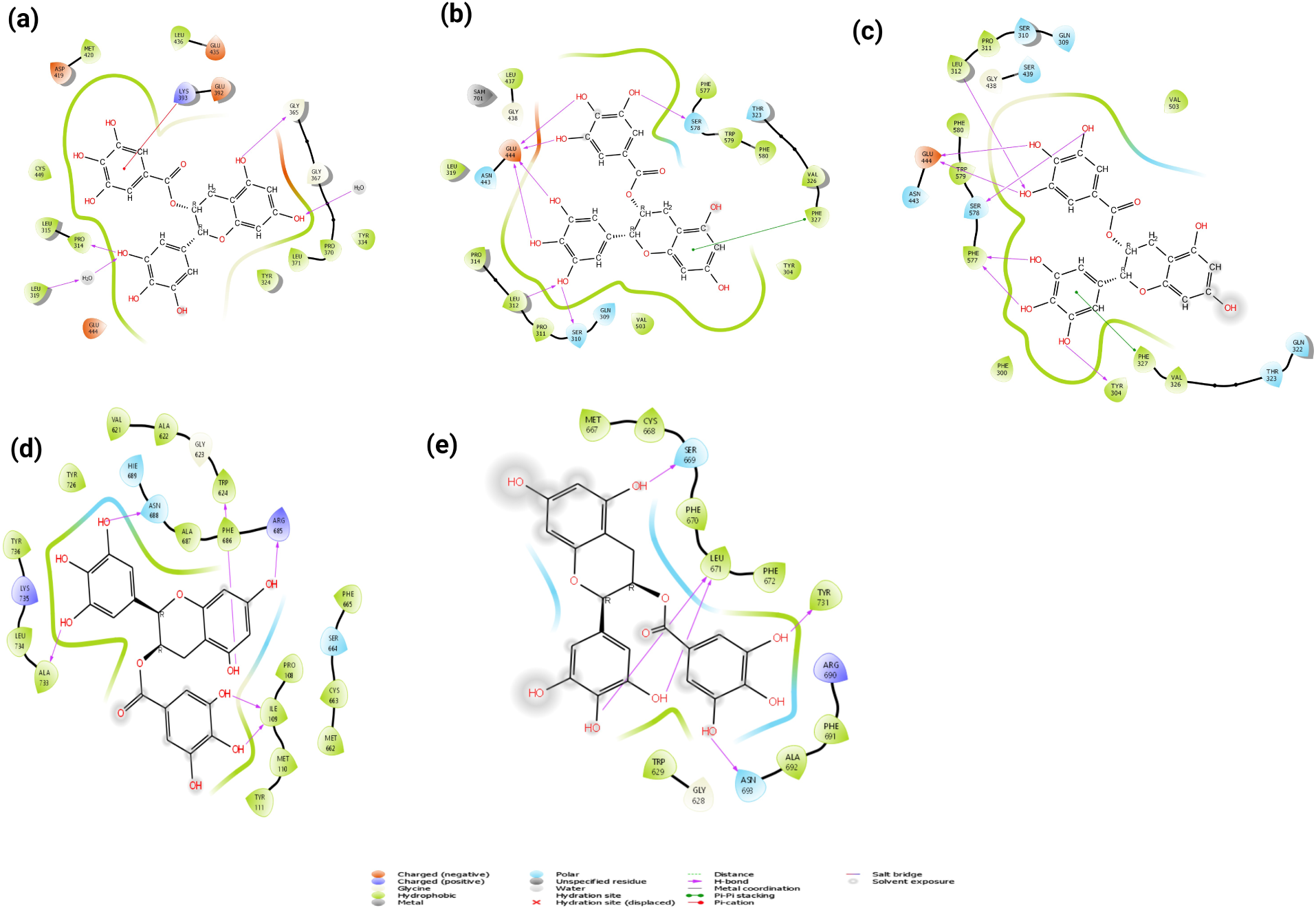
Interaction profile of EGCG with human PRMT5: MEP50 complex. (a-c) Interaction profile of EGCG with PRMT5(4X60, 4X61, 4X63). The hydrogen bonds are shown in green dotted lines, the interacting amino acid residues, as well as the ligand Sfg, are shown in blue/black color ball and stick model, the non-bonded interactions are depicted with residues with starbursts. (d-e) interaction profile of EGCG with EZH2 (5HYN & 4mi5). The ligands are shown in the middle, and the hydrogen bonds are shown with purple arrows, and the arrowheads display the H-bond donor-acceptor relationship. The π-cation interaction of both EGCG and Sfg with the lysine side chain is depicted with a red arrow. The hydrogen bonds are shown in green dotted lines, the interacting amino acid residues, as well as the ligand SFG, are shown in blue/black color ball and stick model, the non-bonded interactions are depicted with residues with starbursts

**Table1:**
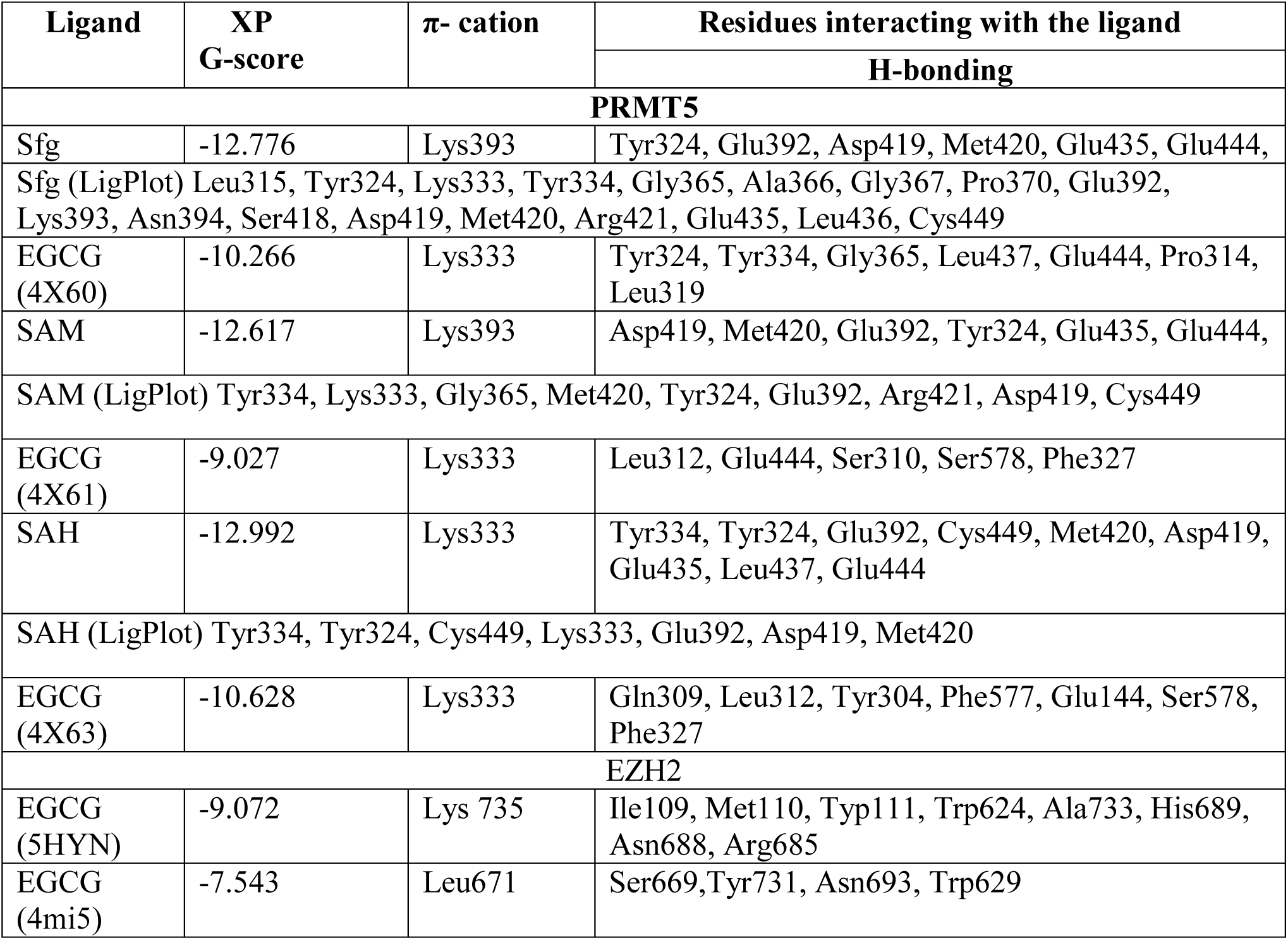
Common interaction profiles of the EGCG with human PRMT5: MEP50 and EZH2: The protein complex (4X60, 4X61, 4X63), & EZH2 (5HYN & 4mi5) were used for the molecular docking studies. The GlideXP docking scores of EGCG, and Sfg, SAM and SAH are also displayed.

### 3.6 Surface Plasmon Resonance studies for EGCG with PRMT5:MEP50 and EZH2

A SPR assay was performed to evaluate the interaction between PRMT5-MEP50, EZH2 with EGCG. The recombinant PRMT5-MEP50 & EZH2 were immobilized on sensor surface of CM5 chip and, sensograms obtained for immobilization is given in (Figure-S2). The interaction and affinity of EGCG with PRMT5-MEP50 & EZH2 was further studied by real time bimolecular interaction analysis. The binding kinetics was studied by running varying concentrations of ligand over immobilized PRMT5-MEP50 and EZH2 surface. (Fig3a&3c). The results indicated that EGCG was shown strong binding affinity in nanomolar concentrations range (200-1.56 nM) with both PRMT5-MEP50 & EZH2 protein. Binding affinity KD (equilibrium constant), was determined to find the strengths of biomolecule interactions. The KD is 1.74 E-05 M and 4.39 E-05 M for EGCG with PRMT5-MEP50 & EZH2 respectively. The graph of response versus concentration is linear (Fig3b-3d) the association constant (ka) and dissociation constant (kd) revealed that EGCG has more affinity for PRMT5 than the EZH2, which is correlating with molecular docking results.

**Figure-3:**
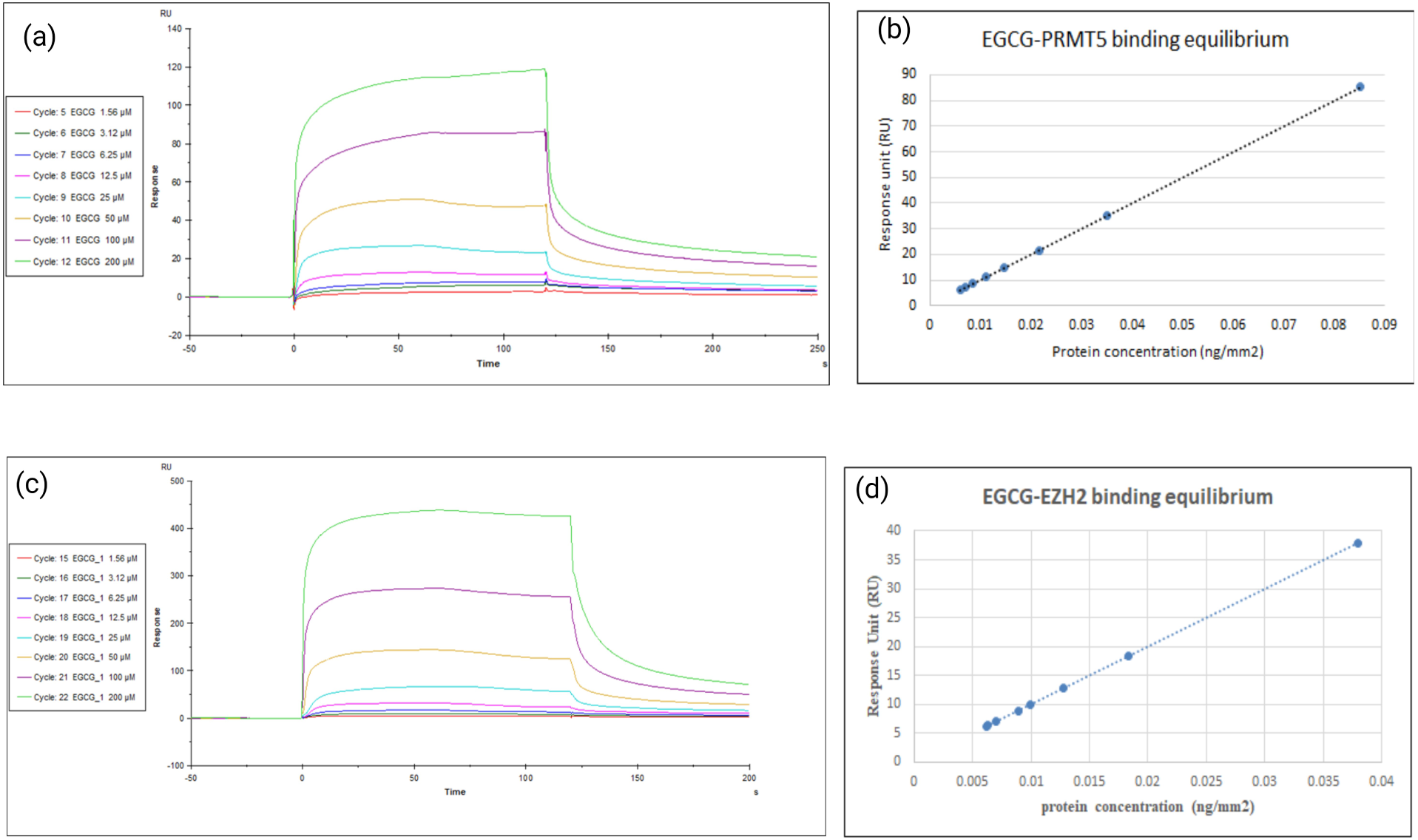
Surface Plasmon resonance Studies: The sensorgram of EGCG on PRMT5 & EZH2 complexes on cm5 sensor chip, (a) dose–response sensorgram of EGCG with immobilized PRMT5-MEP50. (b) Fitting of response and concentration using BIAcore T200 Evaluation software version 2.0. (c) dose–response sensorgram of EGCG with immobilized EZH2. (d) Fitting of response and concentration using BIAcore T200 Evaluation software version 2.0. The response units RU versus protein concentration plot linear. The obtained Ka, kd & KD values are mentioned: for PRMT5 Ka: 2.07E+02, kd 3.61 E-03, & KD-1.74E-05, EZH2: Ka-82.78, kd:3.63 E-03 & KD:4.39E-05 respectively

### 3.7 Impact of EGCG on growth of multiple cancer cell lines

The human cancer cell lines DU-145 (prostate), A549 (lung), HeLa (Cervix), Hep G2 (liver), MCF7 (breast), MDA-MB-231 (breast) PC-3 (prostate) and HEK293 (human embryonic kidney) were included in the study, cells were treated with varying concentrations of EGCG for 24 and 48 hrs and cell viability was accessed to find the IC_50_ value (Fig-S3). The trypan blue dye exclusion assay supported MTT assay and increased concentration of EGCG reduced the viable cell number (Fig4b). As shown in (Fig4a) MDA-MB231 cells incubated with indicated concentration of EGCG for 24 hours and checked the level of catalytic products of PRMT5 & EZH2 i.e., H4R3me2s & H3K27me3, results demonstrated that both H4R3me2s & H3K27me3, levels decreased significantly (particularly at 10 μM concentration) compared to the untreated cells whereas protein level of PRMT5, EZH2, and H4 & H3 levels did not get altered significantly. The β-actin used as a loading control.

**Figure-4:**
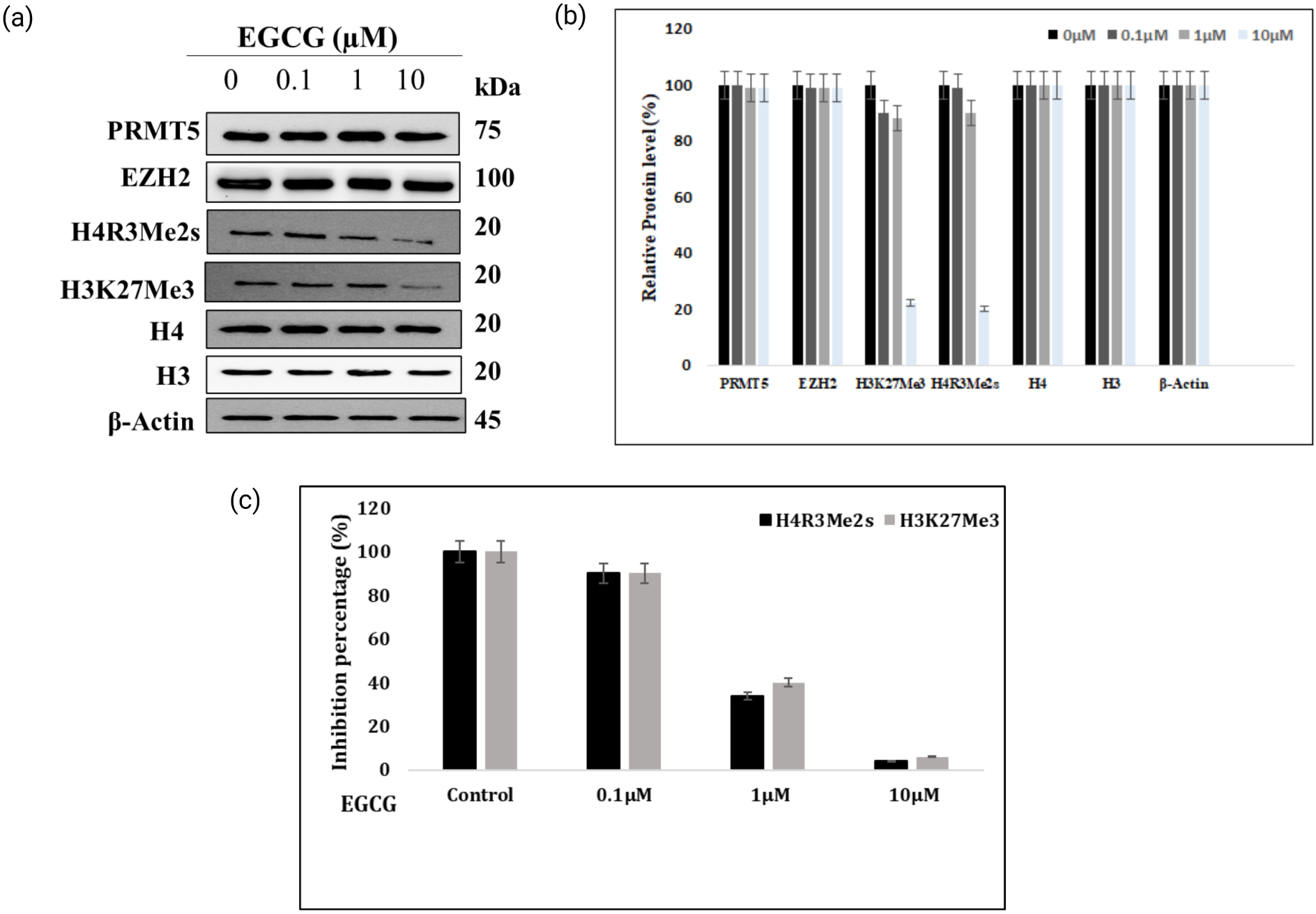
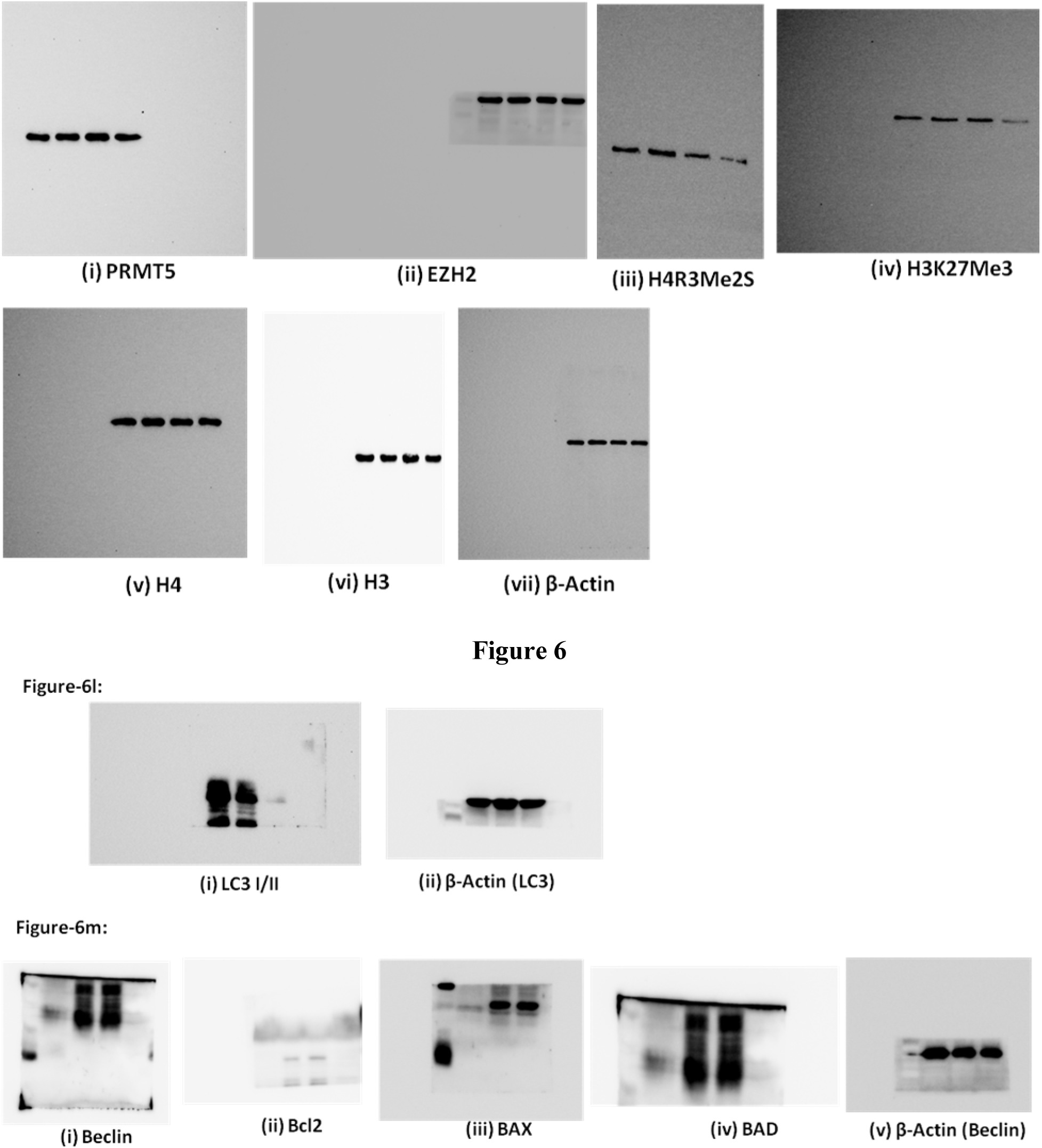
EGCG impact on methylation marks: The MDA-MB231 cells were exposed to increasing concentration of either EGCG (0.1, 1, 10 μM) for 24 hours. Protein lysate was prepared as explained in methods and analyzed for H4R3me2s & H3K27me3 using immunoblotting. The β-actin was used as a loading control. (a) Western blots (b) protein intensity graphs (c) ELISA of *In-vitro* methylation which was carried using EZH2 and PRMT5-MEP50 enzyme complex and histones as a substrate.

The *in*-vitro methylation of H4/H3 was performed using PRMT5-MEP50 and EZH2 and methylation of H3 and H4 was validated through ELISA. As shown in (Fig4c) pre-incubation of PRMT5-MEP50 and EZH2 with EGCG (1 µM, 10 µM) decreased the symmetrical dimethylation of 3^rd^ arginine of histone H4 & trimethylation of lysine 27 on histone H3 in dose dependent manner. These results clearly demonstrated that EGCG strongly inhibiting both PRMT5-MEP50 & EZH2. AO-EtBr double staining, we observed that MDA-MB 231 cells labelled with EtBr/AO after 24 hrs of incubation with EGCG, revealed that the early-stage apoptotic cells appear like crescent shaped or granular ones were observed in EGCG treated samples, while no significant apoptosis was observed in the control group (Fig 5a), which directly correlated by trypan blue dye exclusion assay and MTT assays. (Fig5b). And in the colony formation assay, the EGCG reduced the formation of colonies in dose dependent manner, when compared with control (Fig 5c-5d). The drastic reduction in the formation of colonies observed with increasing concentration of the compound (Fig 5). These invitro assays demonstrated that EGCG potentially inhibit the PRMT5 and EZH2 activity.

**Figure-5:**
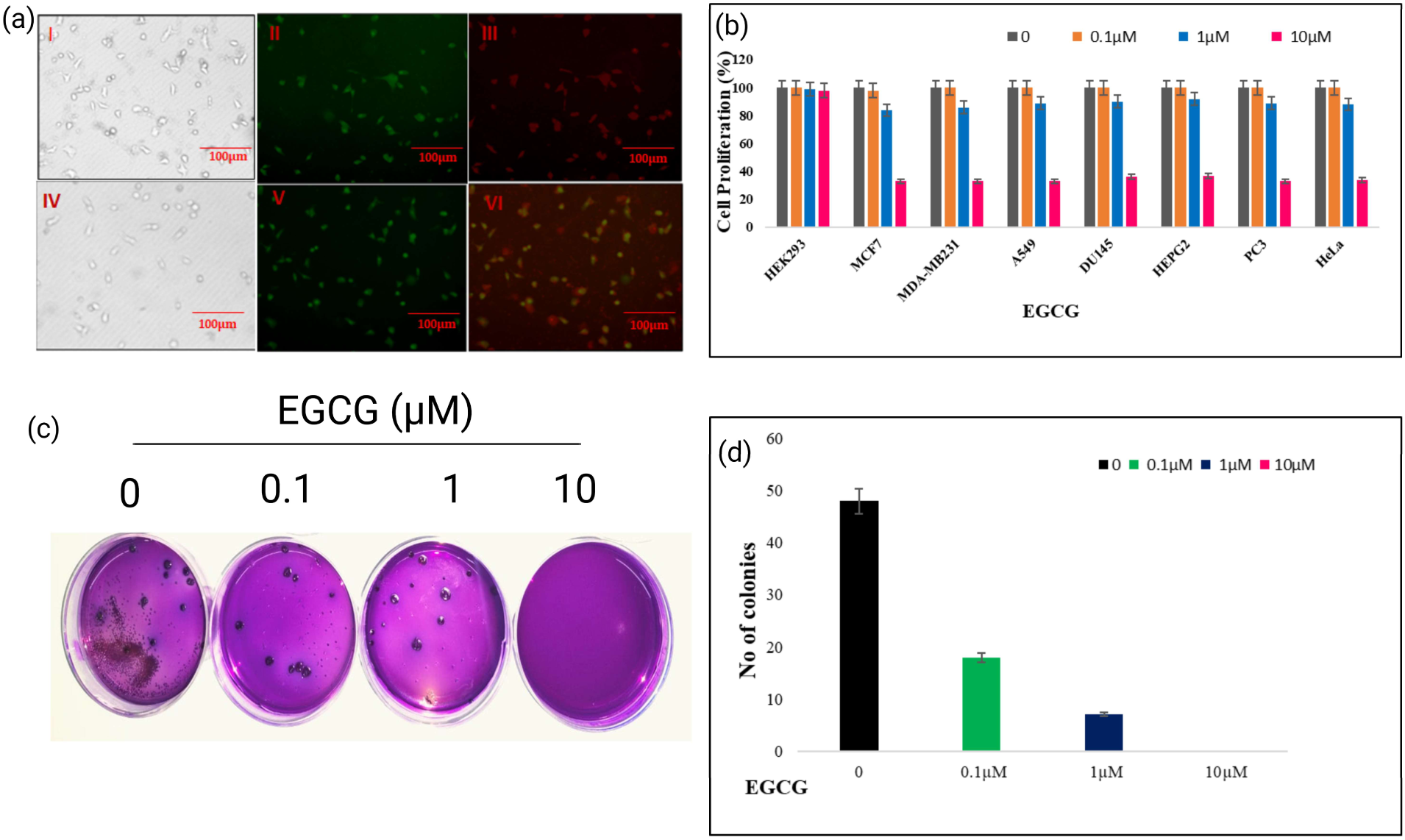
Impact of EGCG on cell proliferation: MDA-MB-231 cells were treated with different concentrations of drug 0, 0.1, 1 and 10 μM respectively, in-vitro assays were performed (a) AO/EtBr assay: Double staining with ethidium bromide and acridine orange (EB/AO) MDA-MB231 cells, treated with EGGC for 24 hours, were observed under a fluorescent microscope, revealing viable cells emitting green fluorescence (AO) and necrotic cells exhibiting dominant red fluorescence (EB). (b) Trypan blue dye exclusion assay : After treatment, Du-145, A549, HeLa, Hep G2, MCF7, MDA-MB-231, PC-3 cells were incubated with Trypan Blue solution, allowing viable cells to exclude the dye, while non-viable cells absorbed the dye, appearing blue. Cell viability was assessed under a microscope, and cells were counted under microscope and results were tabulated and a plotted a graph. (c,d) the colony formation assay was performed, 500 cells seeded were treated as per experimental conditions. Following incubation, colonies were fixed and counted. Comparison between control and treated groups reveals the impact on clonogenic potential, offering insights into altered cell survival and proliferation.

### 3.8 Effect of EGCG on Apoptosis & Cell cycle

Next, we evaluated the effect of EGCG on induction of apoptosis in breast cancer cells. We treated MDA-MB231 cells with increasing concentration (0, 0.1, 1, and 10µM) of EGCG for 24 h, and stained with Annexin V/ PI and determined the percentage of apoptotic cells by flow cytometry. FACS analysis revealed that with gradual increase in concentration of EGCG cells were started entering late apoptosis stage (Fig6a-6d & 6i). Further to examine whether EGCG can affect cell cycle progression, we performed flow cytometry to test cell cycle distribution and evaluated the distribution of cell cycle phases (Fig6e-6h & 6j). Cell cycle analysis showed that EGCG treatment of cells resulted in G0/G1 arrest in a dose-dependent manner at 1 µM and 10 µM compared to untreated control.

**Figure-6:**
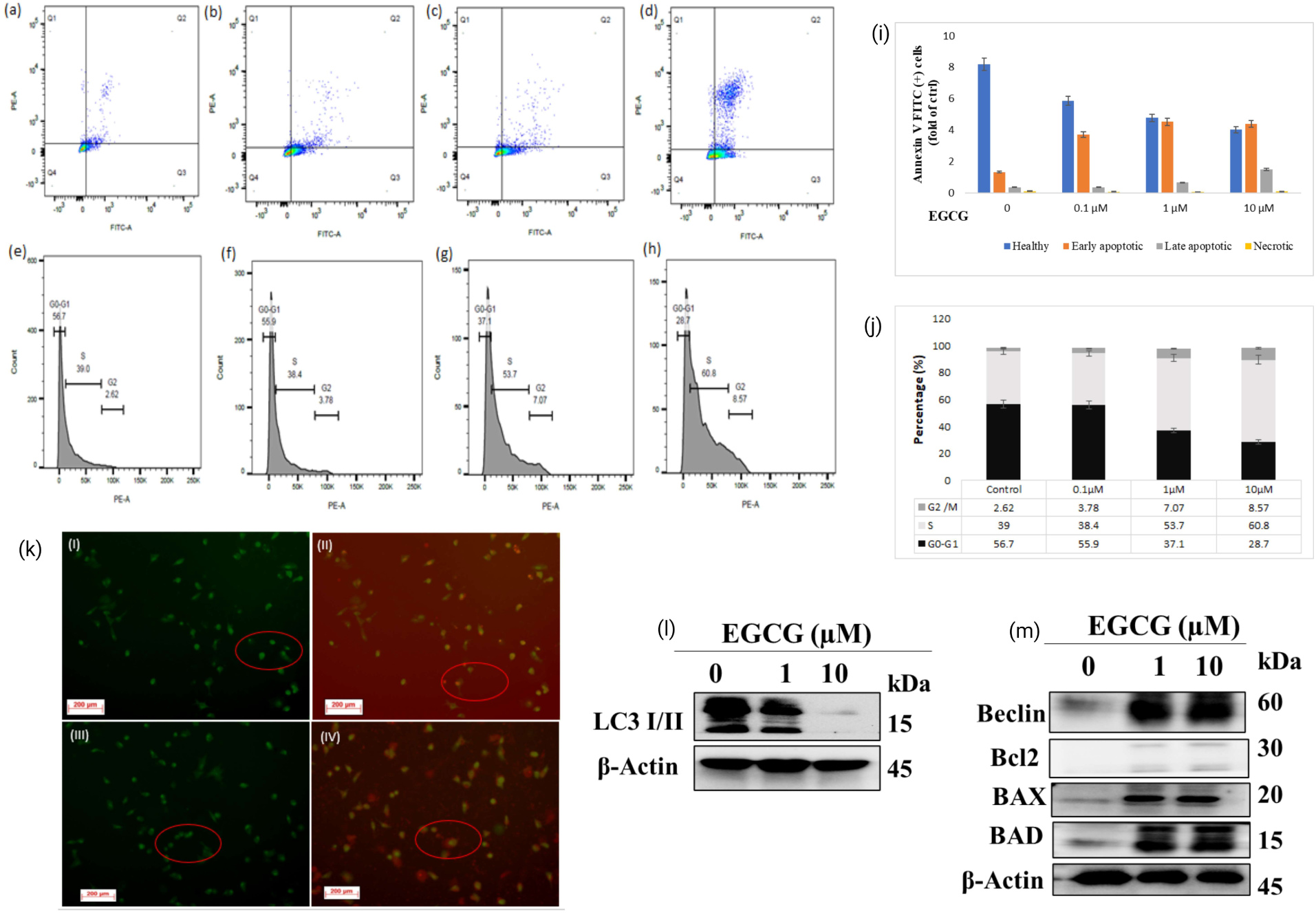
EGCG induces cell death by apoptosis in breast cancer cells: MDA-MB231 cells treated with EGCG for 24 h, and stained with AnnexinV/propidium iodide (PI) and the percentage of apoptotic cells was determined by flow cytometry. Top panel-(a-d) flow cytometry analysis of MDA-MB231 cells upon EGCG treatment, down panel (e-h) cell cycle analysis of MDA-MB231 cells upon EGCG treatment for 24 hrs. EGCG induced autophagy in breast cancer cells (i)-EGCG induced formation of acidic vesicular organelles in MDA MB231 cells (j) western blot analysis to detect LC3 I/II expression in EGCG treated MDA MB231 cells (k) (k)-EGCG induced formation of acidic vesicular organelles in MDA MB231 cells (l) western blot analysis to detect LC3 I/II expression in EGCG treated MDA MB231 cells (m) western blot analysis to check Beclin-1, BCl2, Bax & Bad proteins expression in EGCG treated MDA MB231 cells.

### 3.9 EGCG induced autophagy in MDA-MD 231 cells

AO-EtBr assay showed the appearance of apoptotic cells appear like crescent shaped or granular ones, to reconfirm whether that formation of granular shaped cells whether it is due to apoptosis mediated autophagy we checked for the formation of Acidic vesicular organelles (AVO’s) formation upon EGGC treatment to cells, development of AVO’s is an indicator for autophagy, BY AO (Acridine orange) staining, red fluorescent spots have been observed in EGCG treated cells, whereas control cells showed majorly green cytoplasmic fluorescence (Fig 6k) we further examined in case the EGCG inhibition affected autophagy mediated cell death in MDA-MB231 cells, for this we checked microtubule associated protein light chain 3 (LC3) which is well known marker to monitor the autophagy, Our results also revealed that EGCG caused LC3 transition in a dose-dependent manner (Fig-6l), to ascertain the induction of autophagy, we checked the expression of Beclin-1 and BCL family proteins (BCL2) BAX & BAD (Fig 6m).

### 3.9 EGCG suppressed the tumorigenic potential of MDA-MB231 cells in in-vivo

To determine the anti-tumor effect of EGCG *in-vivo*, MDA-MB231 cells were injected subcutaneously into Swiss nude mice to develop tumor xenografts, and subsequently tumor weight were measured. EGCG (100mg/kg) administration resulted in drastic reductions in tumor volume and compared with those of control groups (**Fig7a-7g)** while the body weight of control group mice increased relatively steady throughout the experiment period. Further to evaluate the potential toxicity of EGCG, H&E staining was performed to tumor tissues, comparison of the control tumors with EGCG treated tumors revealed that in control group observed that highly aggressive sarcomatous neoplastic cells appeared vacuolated appearance with a greater number of mitotic (**Fig7a-7d)** while in treated tumors observed that section has only sub cutaneous and muscular layer, but no dermal and epidermal tumour region **(8a-8d),** and furthermore, the proliferation of cells was by identified using Ki-67 proliferation marker, immunohistochemistry (IHC) revealed that EGCG reduced Ki-67 levels considerably in the tumors treated with EGCG and compared with control group. (**Fig8e-8j).** To reconfirm that EGCG is reducing the catalytic products of both PRMT5 and EZH2, we performed western blot analysis for EGCG treated and control group tumors and observed that of EGCG treated mice tumors showed dramatic reduction of the catalytic products of PRMT5 & EZH2 i.e., H4R3me2s & H3K27me3 respectively **(Fig7e-7f).**

**Figure-7:**
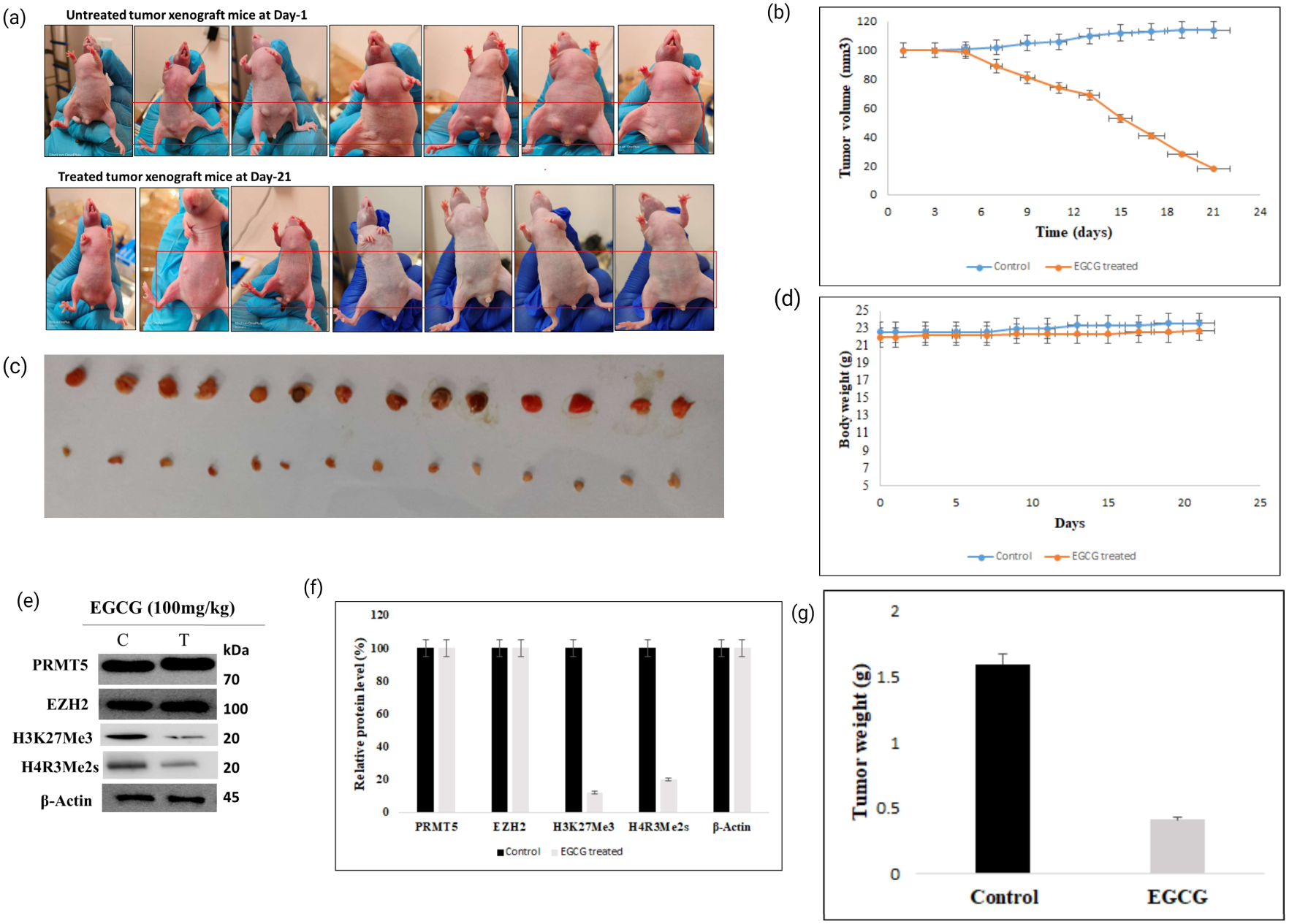
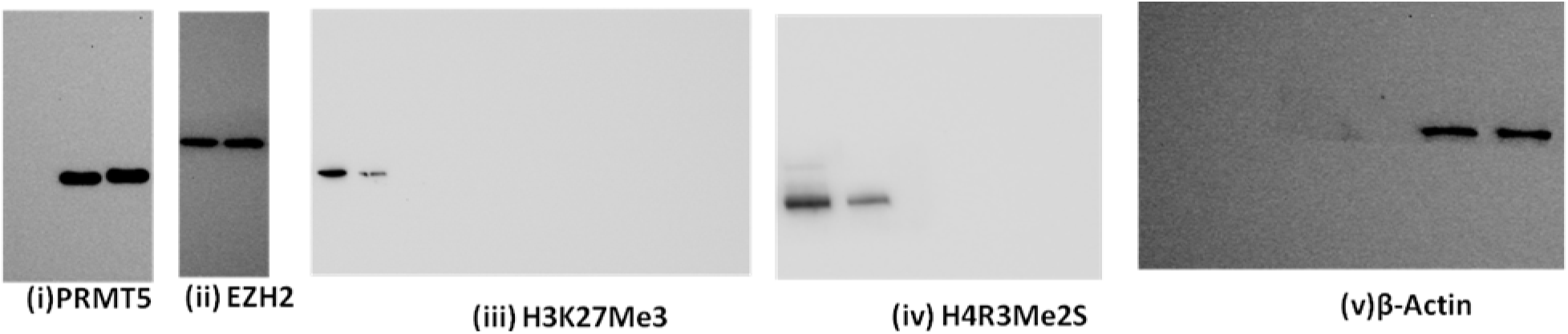
In vivo xenograft studies: The mice were divided into three groups at randomly with 7 mice per group: (1) control group-1 (2) control group-2 (3) treatment group. The mice were injected MDA-MB-231 cells subcutaneously on both flaks. Tumor volume was measured every three days. The tumor volume once reached 100mm^3^ started the EGCG (100mg/kg) administration orally to treatment group for a period of 21 days. (a) comparison of EGCG treated *vs* control from left to right 1^st^ is control mice, remaining are from 2^nd^ to 7^th^ bearing tumors at day 1^st^ of treatment (b) tumor growth measurement top panel in control animals down panel is of treatment (c) Tumor volume comparison graphs (control *vs* treated mice). (d) Body weight measurement graphs (control *vs* treated). (e-f) The western blot analysis of tumor lysates from treatment group, levels of PRMT5 & EZH2 and H3R3mes and H3K27me3, β-actin was used as loading control.. (g) tumor weight comparison graphs (control vs treated)

**Figure-8:**
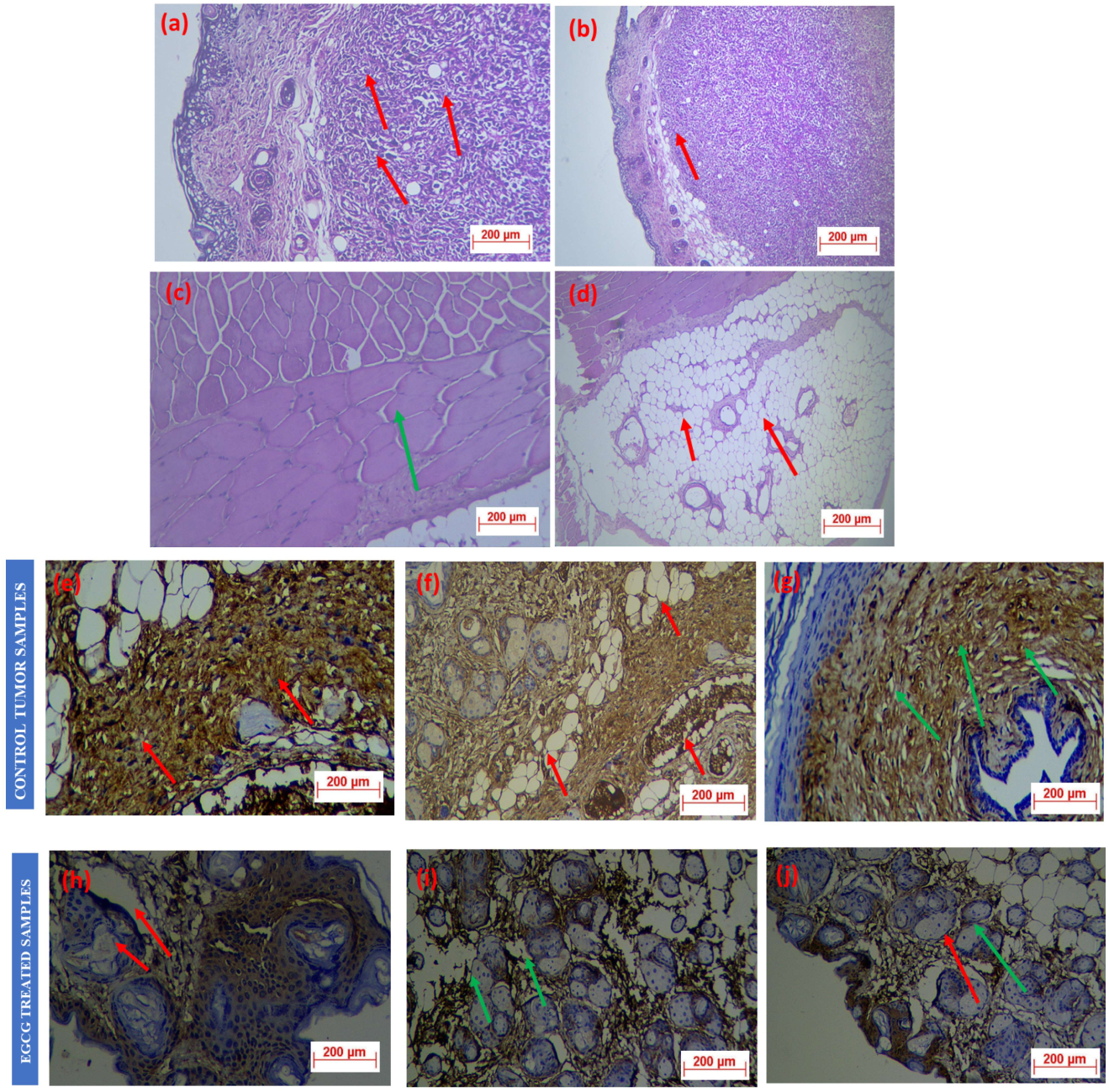
Immunohistochemistry studies: After 21days of drug dosage, mice were euthanized and tumors were collected from each group (control and treated), for IHC studies tumor samples submitted to histopathology expert. Comparison of the control tumors with EGCG drug treated tumors. (8a-8b) H& E staining, in the control samples(tumor) showed highly aggressive neoplastic sarcomatous cells formed a nodule in sub cutaneous region and invaded in to dermal region with mitotic figure indicated in red arrows. (8c-8d) EGCG treated tumor samples showed only sub cutaneous and muscular layer, but no dermal and epidermal tumour region. Subcutaneous region with adipocytes appeared normal and no metastatic invasion of neoplastic cells indicated by red arrow in (8d), and (8c) the muscular region appeared normal and no metastatic invasion of neoplastic cells indicated marked by green arrow. (8e-8g) shows the Ki67 expression, control tumor samples, showed severe expression of Ki 67 in subcutaneous tumour mass indicated by red arrows in 8e, 8f and neoplastic cells invaded in to dermal and dermal hair follicles showed expression indicated by green arrow (8g). The EGCG treated tumor samples (8h-8j) showed that 10-20 percent expression of Ki 67 was noticed in pleomorphic anaplastic epithelial cells in the epidermal layer indicated by red arrows. The Ki67 over expression in stromal tissue in dermal and sub cutaneous region indicated by green arrows (8i,j).

## Discussion

The clinically relevant inhibitors of epigenetic regulators like histone deacetylases (HDAC)s **^(^**^28, 29^**^)^**and DNA methyltransferases (DNMT) **^(30)^**, Histone Methyltransferases (HMTs) **^(31)^**, Histone Demethylase (HDMs) **^(32)^**, Bromodomain and Extra-Terminal (BET) **^(33)^** are well documented. The DNMT inhibitor (DNMTi) and HDAC inhibitor (HDACi) and BET inhibitors (BETi) has already been approved as drugs against selective cancers and entered market for clinical use **^(34)^**. Plethora of small molecules has been targeted with a wide range of potential to inhibit PRMT5 and EZH2. The surge in the increased number of approvals for PRMT5 and EZH2 inhibitors attributed the significant promise in cancer therapeutics **^(^**^10,14, 35,36,37,38^**^).^** Array of data suggested that both PRMT5 and EZH2 is over expressed in several cancers including breast, prostate endometrial, liver, ovarian, small cell lung cancer, melanoma, glioblastoma, and pediatric glioma bladder as well as lymphomas, with positive correlation on disease progression and poor prognosis **^(8,^**^10,39^ ^6,8,35,36,37,38^**^),^** and PRMT5 and EZH2 functionally associate, PRMT5-mediated histone marks, may lead a synergic effect **^(43)^** and hence combined therapeutic approach is promised strategy for therapy. An array of phytocompounds has been reported for the inhibition or modulation of epigenetic regulatory proteins such as DMNTs, HDACs, PRMTs and PRC proteins. The phytocompounds like curcumin, resveratrol, brazilin, catechin, quercetin, and EGCG etc are shown to be physically interacting with several proteins including the DNMT1, HDAC1 in its catalytic pocket **^(34)^**. The phytochemicals have a tremendous impact on epigenetic regulators leading changes via the direct or indirect effect on activity and showed promising role in cancer prevention and therapy. For instance, genistein, curcumin, resveratrol reduces DNMT activity and resulting in activation of tumor suppressor genes, sulforaphane, forms complex with the active sites of HDACs thereby impeding HDAC activity **^(44)^**.

A large number of studies in animals, as well as epidemiologic, case-control studies in humans, reveal that EGCG significantly reduce the incidence of various cancers **^(44)^**. Its therapeutic action was observed to inhibit multiple enzymes, including DNA topoisomerase, tyrosinase, arylamine N-acetyltransferase, glycogen phosphorylase, glutathione transferase, inositol-trisphosphate 3-kinase, inositol-polyphosphate multi kinase, nucleoside-diphosphate kinase, and DNA-directed DNA polymerase **^(17,18,19,44)^**. It is also reported that it induces epigenetic modifications, angiogenesis, carcinogenesis, metastasis and enhance the efficacy of conventional anticancer drugs also **^(19)^**. In present study we first time report the inhibition of PRMT5 and EZH2 activity by EGCG. The molecular docking studies revealed that the EGCG forms a cation-π non-covalent interaction between the positively charged lysine (333) side chain and the phenolic ring of EGCG containing π electrons; while at least five hydrogen-binds were formed involving the key residues within the catalytic pocket of PRMT5 residues Tyr324, Tyr334, Gly365, Leu437, and Glu444 within the SAM-dependent methyltransferase of double E loop. EGCG forms an important π-π stacking with Phe327 and more interestingly Glue435 residue involved in the interaction. The binding site is sharing common residues with sinefungin and in close proximity to the SAM-binding region, which signifies the catalytic inhibitory potential of EGCG that might have contributed to the reduced formation of H4R3me2s methylation marks in the EGCG-exposed cells. As previous studies reported, **^(21,45)^** it requires the water molecule to mediate interaction with pi-pi stacking interaction phenolic ring. Notably, this may be due to conformational changes occurs while placing SAM analogue. This observation is well coinciding with SPR binding studies where strong binding affinity for EGCG at (1.74 E-05 M (PRMT5:MEP50) and 4.39 E-05 M (EZH2)). Interestingly, when compare to previous studies the binding interactions of most well studied synthetic molecule EPZ015666 binds to PRMT5 independent from SAM binding sites and other molecule PR5-LL-CM01 binding interactions span over the Rossman fold and β-barrel domains of PRMT5, provided that EGCG more potent than EPZ015666 and PR5-LL-CM01**^(46)^**. While the EZH2 (5HYN), interacted with five hydrogen bonds (Trp624, Ile109, Ala733, Asn688 and His689), with 4mi5 Leu671, Asn698, Ser669 and Tyr731. The study revealed that EGCG is more potently binding to PRMT5 than the EZH2 which has been further evidenced with surface plasma resonance binding assay. The *In vitro* methylation assays followed by ELISA and reduction in the level specific histone marks upon treatment in cells strongly supported the inhibition of activity by EGCG in MCF-7 and MDA-MB231 cells.

The inhibition of activity or down regulation of both PRMT5/EZH2 and its effects on multiple process like induction of apoptosis, autophagy and cell cycle arrest in multiple cancer studies has been reported **^(17,18,19)^**. It has been observed that EGCG caused caspase-3 activation and increased the expression of Bax in PEL cells, indicated the EGCG mediated induction of apoptosis **^(62)^**. Meantime, its treatment also increased the acidic vesicular organelles formation and the expression of LC3-I/II and beclin-1, divulged that EGCG can also induce autophagy **^(^**^62–63^**^)^**. Autophagy plays dual roles in cancer; it can help the survival of cancer cells as well as promote cell death. Taking the evidence of the inhibition of EZH2 by a specific inhibitor, 3-deazaneplanocin A (DZNep), caused the apoptosis of NRK-52E cells via modulating H3K27me3 in its promoter region it suggested that reduced activity of EZH2 by EGCG may cause the autophagy as observed in the present study. In another study showed that downregulation of EZH2 is critical for the induction of autophagy and apoptosis in colorectal cancer cells and EZH2-siRNA inhibits the proliferation and promoting apoptosis by inducing G0/G1 cell cycle arrest of SW620 cells. The level of histone H3 lysine 27 trimethylation, altering cell cycle regulator cyclins and p21 an p27 in skin cancer **^(49)^** substantiate the EGCG mediated reduction of H3K27me3. The GSK591 induced inhibition of PRMT5 or shRNAs mediated down regulation induce the apoptosis and autophagy, with reduced Akt/GSK3β phosphorylation targets like cyclin D1 and E1 in human lung cancer cell death **^(50)^**. The knockdown of PRMT5 expression enhanced cell pyroptosis in multiple myeloma **^(51)^** and PRMT5 regulates expression of cell cycle/apoptosis-associated genes by modifying p53, E2F-1 and KLF4 **^(10)^**. PRMT5 inhibition in high-risk multiple myeloma leads to increased apoptosis and decreased cell growth **^(52)^**. PRMT5 activity has effect on different stages of autophagy thereby control autophagy and tumorigenesis through modulating the functions of specific targets **^(53)^**. In triple negative breast cancer cells, PRMT5 catalyzes ULK1 monomethylation at R532 to suppress ULK1 activation, and have a role in autophagy **^(54)^**. Trichostatin A mediated suppresses of cervical cancer cell proliferation and induces apoptosis and autophagy via regulation of the PRMT5/STC1/TRPV6/JNK axis **^(55)^**.Thus the global loss of the repressive histone methylation marks, like H4R3me2s, and H3K27me3 may cause the induction of apoptosis and autophagy reported in the present study, further these marks also known to be associated cell cycle arrest, we hypothesized that EGCG mediated inhibition is responsible to arrest of cell cycle, in the current study which is evidenced with earlier reports the where EGCG arrested cell cycle at G0/G1 phase of cell cycle **^(17,53,54,55)^** G2/S**^(17)^** and/or G2/M **^(17,53,54)^** cell cycle in variety of cancer models. This event is through a different mediator like cyclin D1, cdk4, cdk6, p21 and p27 in pancreatic cancer cells **^(56)^** and via EGFR/cyclinD1signaling in A549, H460 and H1650 lung cancer cells **^(57)^** and by WAF1/p21-in LNCaP and DU-145 prostate cancer cells **^(54)^**. As PRMT5 has many non-histone substrates that involved in cell cycle regulation and cell proliferation **^(10)^** thus the repressive histone marks may directly or indirectly involve in modulating functions of these proteins.

Many studies using cell lines and animal models have demonstrated anticancer activity of EGCG **^(16–19,47,48,57)^**. The information from clinical trials suggested a safety of EGCG (200 mg/day) in men for prostate cancer **^(58)^**. In a randomized another report, phase II pilot study in bladder cancer patients, showed an EGCG accumulation in cancer tissue with decreased the level of proliferation **^(59)^**. In a phase 1 trial showed the chemo preventive effects of Teavigo™ (highly purified and refined green tea extract providing 94% EGCG) (450 mg/PO/day) in colorectal cancer (CRC) patients (NCT03072992). **^(60)^**. In our study also observed the drastic reduction in tumor size with reduced level of histone marks in MDA-MB231 induced xenografts which is supported by massive reduction of proliferative marker Ki76 even though the direct link between Ki67 and reduced level of H4R3mes and H3K27me3 lacking. The inhibitory impact of EGCG on the catalytic activity of the oncogenic PRMT5 and EZH2 could serve as a potent anti-cancer modality and natural molecule further might be established as potent PRMT5/EZH2 inhibitor.

Finally taken together, the results of current study describe the once again potential of EGCG which is already known for their anti-cancer effects and in clinical trials **^(16–19)^** as a novel interacting molecule to the most sought-after methyltransferase PRMT5 and PRC2 complex-histone methyltransferase EZH2. The results demonstrated *in-silico*, *in-vitro* studies and SPR binding studies, which potentially inhibit activity with diminished the level of the repressive histone methylation marks. Furthermore *in vivo* studies revealed that oral dosage significantly reduced tumor size, and proliferation marker like ki67. The EGCG mediated inhibition of both PRMT5 and EZH2 can potentially to be developed as novel strategy to develop drug.

## Abbreviations

PRMT5: protein arginine methyltransferase 5
EZH2: enhancer of Zeste homolog 2
PRC2: Polycomb Repressive Complex 2
MEP50: methylosome protein 50
PRMTs: Protein arginine methyltransferases;
H3K27me3: trimethylated lysine 27 on histone H3
H4R3me2s: symmetrically dimethylated arginine 4 on histone H4
H4R3: H4 residue on Arg3
H3K27: H3 residue on lys27, EGCG, Epigallocatechin-3-gallate
SPR: surface plasmon resonance
MMA: monomethylarginine
ADMA: asymmetric dimethylarginine
SDMA: symmetric dimethylarginine, TCGA, the Cancer Genome Atlas
SMILES: Simplified molecular-input line-entry system
SAM: S-adenosyl methionine
AVO: acidic vesicular organelles, MTT, 3-(4,5-Dimethylthiazol-2-yl)-2,5Diphenyltetrazolium-Bromide.

## Author contributions

NKK: Performed the experiments and wrote the manuscript; BC: Performed the initial screening of compounds. JP; Helped and performed molecular docking, KG: Conceptualized the *in silco* work AS: Helped in Insilco and molecular docking studies AP; Provided molecular docking platform; CJ; Provided the molecular docking software platform and critical comments, BM: Helped in animal studies for in vivo studies, and critical comments. SRK: Conceptualized,

designed the experiments, interpreted the data, supervised the entire work, and wrote the manuscript.

## Conflict of Interest

None

## Acknowledgement

We acknowledge the financial support of Institute of Eminence (IoE) (UoH-IoE-RC3-21-008) by University of Hyderabad, and Council of Scientific and Industrial Research, Govt. of India (Grant No. 38(1481)/19/EMR-II) and acknowledged DST-FIST, UGCSAP, DBT-BUILDER infrastructural facilities of the department. NKK thankful to UoH-Non-Net fellowship and DST-INSPIRE fellowship (DST/INSPIRE/03/2017/000035 & IVR No-201700011820 dated 4^th^ July 2018) and AS DST-INSPIRE fellowship (DST/INSPIRE/03/2018/000039 and IVR No.: 201800024300)”.provided by Department of Science and Technology, Govt. of India. We are highly grateful to Mrs. Monika Kannan (Proteomics Facility, School of Life Sciences, UoH) and Dr K Praveen (Cytiva) for helping in the analysis of SPR data.

## Supplementary figures

**Figure S1:**
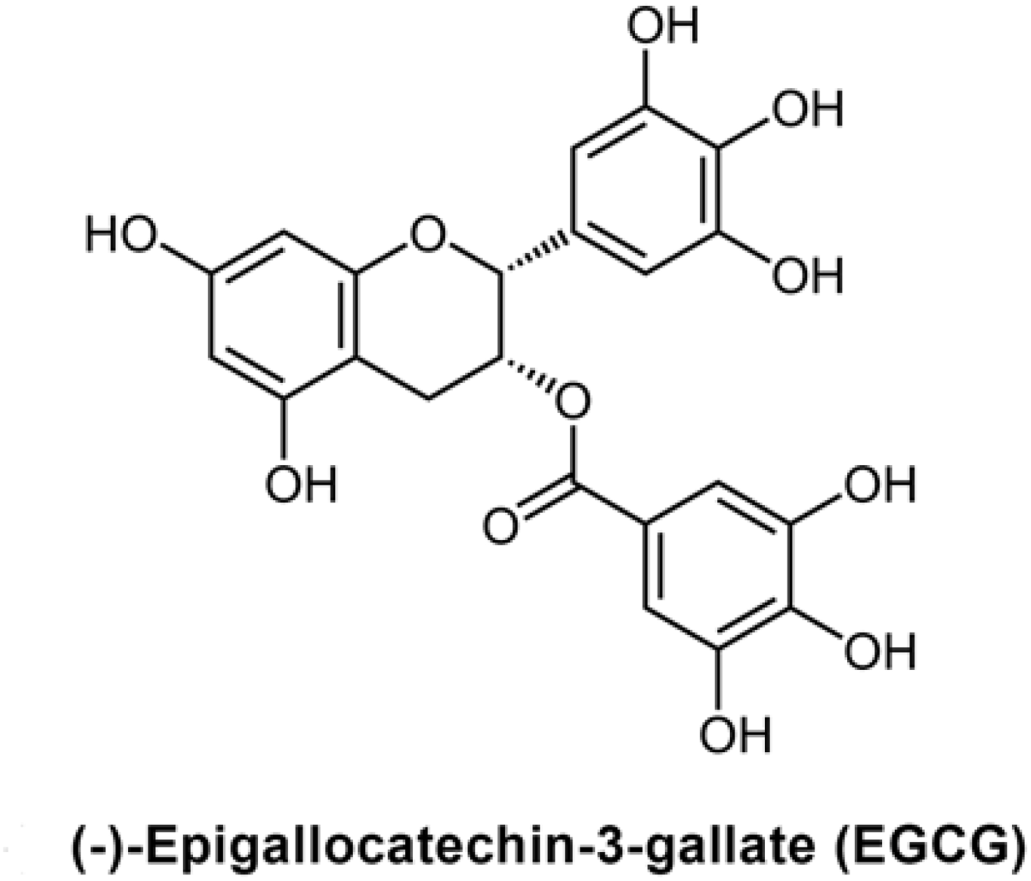
Chemical structure of Epigallocatechin-3-gallate (EGCG) (Mol wt. 458.372 g/mol, chemical formula: C_22_H_18_O_11_

**Figure S2:**
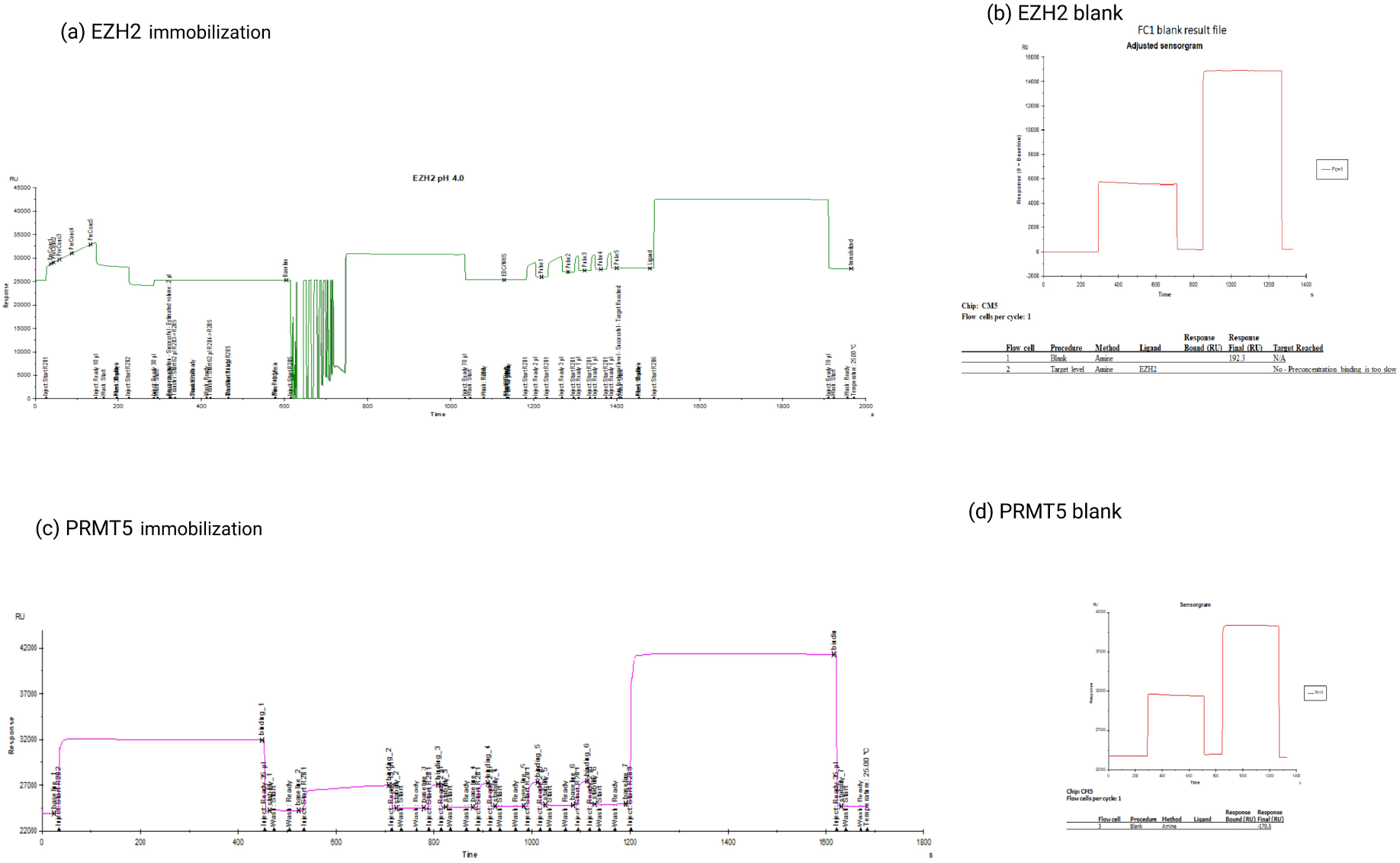
SPR Stenographs of immobilized PRMT5 & EZH2 on CM5 chip. (a &c): PRMT5 & PRMT5 immobilization on CM5 sensor chip (b & d): EZH2 & PRMT5 blanks

**Figure-S3:**
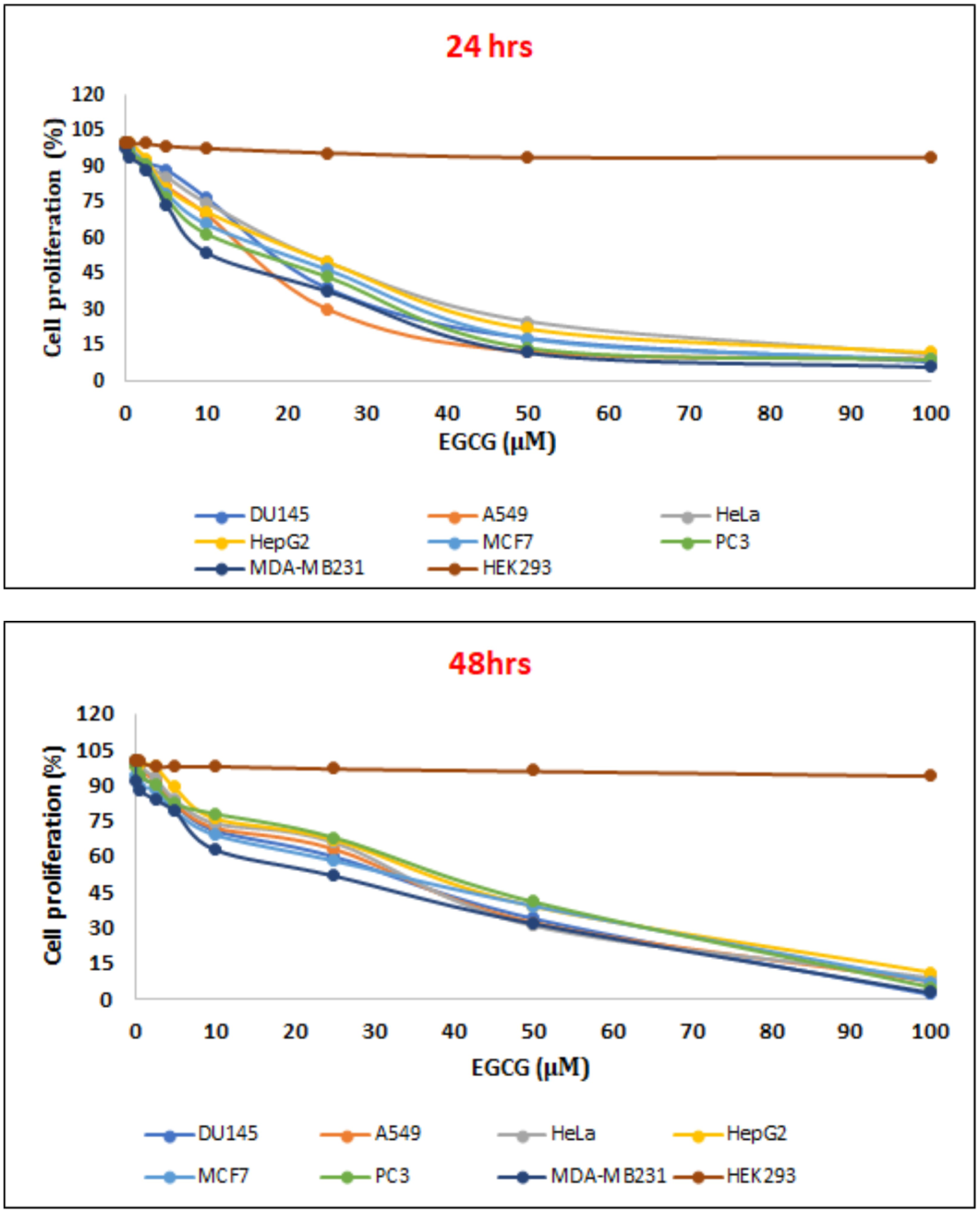
Impact of EGCG on human malignant cell lines: The human cancer cell lines Du-145 (prostate), A549 (lung), HeLa (Cervix), Hep G2 (liver), MCF7 (breast), MDA-MB-231 (breast) PC-3 (prostate) and HEK293 (human embryonic kidney) were exposed to different concentrations of EGCG for 24 & 48 hours for assessing the cyto-toxicity of EGCG and calculated Ic50 values and tabulated.

## Notes

### Competing Interest Statement

The authors have declared no competing interest.

## References

1. Siegel, R. L., Miller, K. D., Wagle, N. S. & Jemal, A. (2023). Cancer statistics, 2023. CA: a cancer journal for clinicians, 73(1), 17–48. 10.3322/caac.21763

2. Sung H, Ferlay J, Siegel RL, Laversanne M, Soerjomataram I, Jemal A, Bray F. Global Cancer Statistics 2020: GLOBOCAN Estimates of Incidence and Mortality Worldwide for 36 Cancers in 185 Countries. CA Cancer J Clin. 2021 May;71(3):209–249. doi: 10.3322/caac.21660. Epub 2021 Feb 4. PMID: 33538338.

3. Hoxha, I., Sadiku, F., Hoxha, L., Nasim, M., Christine Buteau, M. A., Grezda, K., & Chamberlin, M. D. (2024). Breast Cancer and Lifestyle Factors: Umbrella Review. Hematology/oncology clinics of North America, 38(1), 137–170. 10.1016/j.hoc.2023.07.005

4. Li, D., Peng, X., Hu, Z., Li, S., Chen, J., & Pan, W. (2024). Small molecules targeting selected histone methyltransferases (HMTs) for cancer treatment: Current progress and novel strategies. European journal of medicinal chemistry, 264, 115982. 10.1016/j.ejmech.2023.115982

5. Wu, K., Niu, C., Liu, H., & Fu, L. (2023). Research progress on PRMTs involved in epigenetic modification and tumour signalling pathway regulation (Review). International journal of oncology, 62(5), 62. 10.3892/ijo.2023.5510

6. Huang, J., Gou, H., Yao, J., Yi, K., Jin, Z., Matsuoka, M., & Zhao, T. (2021). The noncanonical role of EZH2 in cancer. Cancer science, 112(4), 1376–1382. 10.1111/cas.14840

7. Halabelian, L., & Barsyte-Lovejoy, D. (2021). Structure and Function of Protein Arginine Methyltransferase PRMT7. Life (Basel, Switzerland), 11(8), 768. 10.3390/life11080768

8. Liu, Y., & Yang, Q. (2023). The roles of EZH2 in cancer and its inhibitors. Medical oncology (Northwood, London, England), 40(6), 167. 10.1007/s12032-023-02025-6

9. Yang, X., Xu, L., & Yang, L. (2023). Recent advances in EZH2-based dual inhibitors in the treatment of cancers. European journal of medicinal chemistry, 256, 115461. 10.1016/j.ejmech.2023.115461

10. Kim, H., & Ronai, Z. A. (2020). PRMT5 function and targeting in cancer. Cell stress, 4(8), 199–215. 10.15698/cst2020.08.228

11. Antonysamy Stephen, Zahid Bonday, Robert M Campbell, Brandon Doyle, Zhanna Druzina, Tarun Gheyi, Bomie Han, Louis N Jungheim, Yuewei Qian, Charles Rauch, Marijane Russell, J Michael Sauder, Stephen R Wasserman, Kenneth Weichert, Francis S Willard, Aiping Zhang, Spencer Emtage Crystal structure of the human PRMT5:MEP50 complex Proc Natl Acad Sci U S A 2012 Oct 30;109(44):17960–5

12. Webb LM, Sengupta S, Edell C, Piedra-Quintero ZL, Amici SA, Miranda JN, Bevins M, Kennemer A, Laliotis G, Tsichlis PN & Guerau-de-Arellano M (2020) Protein arginine methyltransferase 5 promotes cholesterol biosynthesis–mediated Th17 responses and autoimmunity. J Clin Invest 130, 1683–1698.

13. Motolani, A., Martin, M., Sun, M., & Lu, T. (2021). The Structure and Functions of PRMT5 in Human Diseases. Life (Basel, Switzerland), 11(10), 1074. 10.3390/life11101074

14. Jarrold J, Davies CC. PRMTs and Arginine Methylation: Cancer’s Best-Kept Secret? Trends Mol Med. 2019 Nov;25(11):993–1009. doi: 10.1016/j.molmed.2019.05.007. Epub 2019 Jun 20. PMID: 31230909.

15. Palomba, M. L., Cartron, G., Popplewell, L., Ribrag, V., Westin, J., Huw, L. Y., Agarwal, S., Shivhare, M., Hong, W. J., Raval, A., Chang, A. C., Penuel, E., & Morschhauser, F. (2022). Combination of Atezolizumab and Tazemetostat in Patients with Relapsed/Refractory Diffuse Large B-Cell Lymphoma: Results from a Phase Ib Study. Clinical lymphoma, myeloma & leukemia, 22(7), 504–512. 10.1016/j.clml.2021.12.014

16. Wang, L., Li, P., & Feng, K. (2023). EGCG adjuvant chemotherapy: Current status and future perspectives. European journal of medicinal chemistry, 250, 115197. 10.1016/j.ejmech.2023.115197

17. Ferrari, E., Bettuzzi, S., & Naponelli, V. (2022). The Potential of Epigallocatechin Gallate (EGCG) in Targeting Autophagy for Cancer Treatment: A Narrative Review. International journal of molecular sciences, 23(11), 6075. 10.3390/ijms23116075

18. Alam, M., Ali, S., Ashraf, G. M., Bilgrami, A. L., Yadav, D. K., & Hassan, M. I. (2022). Epigallocatechin 3-gallate: From green tea to cancer therapeutics. Food chemistry, 379, 132135. 10.1016/j.foodchem.2022.132135

19. Kciuk, M., Alam, M., Ali, N., Rashid, S., Głowacka, P., Sundaraj, R., Celik, I., Yahya, E. B., Dubey, A., Zerroug, E., & Kontek, R. (2023). Epigallocatechin-3-Gallate Therapeutic Potential in Cancer: Mechanism of Action and Clinical Implications. Molecules (Basel, Switzerland), 28(13), 5246. 10.3390/molecules28135246

20. Della Via, F. I., Alvarez, M. C., Basting, R. T., & Saad, S. T. O. (2023). The Effects of Green Tea Catechins in Hematological Malignancies. Pharmaceuticals (Basel, Switzerland), 16(7), 1021. 10.3390/ph16071021

21. Friesner, R. A.; Murphy, R. B.; Repasky, M. P.; Frye, L. L.; Greenwood, J. R.; Halgren,T. A.; Sanschagrin, P. C.; Mainz, D. T., “Extra Precision Glide: Docking and Scoring Incorporating a Model of Hydrophobic Enclosure for Protein-Ligand Complexes,” J. Med. Chem., 2006, 49, 6177–6196

22. Rao, K. S., Nalla, K., Ramachandraiah, C., Chandrasekhar, K. B., Kanade, S. R., & Saha, S. (2023). Design and Synthesis of New Triazole-Benzimidazole Derivatives as Potential PRMT5 Inhibitors. ChemistrySelect, 8(11), e202204474.

23. Franken, N. A., Rodermond, H. M., Stap, J., Haveman, J. & van Bree, C. (2006). Clonogenic assay of cells in vitro. Nature protocols, 1(5), 2315–2319. 10.1038/nprot.2006.339

24. Laemmli, U. K. (1970). Cleavage of structural proteins during the assembly of the head of bacteriophage T4. Nature 227(5259): 680–685

25. Cheng D, Vemulapalli V, Bedford MT (2012) Methods Applied to the Study of Protein Arginine Methylation. Methods in Enzymology 512:71–92.

26. Lim, H. K., Lee, H., Moon, A., Kang, K. T., & Jung, J. (2018). Exploring protocol for breast cancer xenograft model using endothelial colony-forming cells. Translational Cancer Research, 7(5), 1228–1234.

27. Fischer, A. H., Jacobson, K. A., Rose, J., & Zeller, R. (2008). Hematoxylin and eosin staining of tissue and cell sections. CSH protocols, 2008, pdb. prot4986. 10.1101/pdb.prot4986

28. Ramaiah, M. J., Tangutur, A. D., & Manyam, R. R. (2021). Epigenetic modulation and understanding of HDAC inhibitors in cancer therapy. Life sciences, 277, 119504. 10.1016/j.lfs.2021.119504

29. McClure, J. J., Li, X., & Chou, C. J. (2018). Advances and Challenges of HDAC Inhibitors in Cancer Therapeutics. Advances in cancer research, 138, 183–211. 10.1016/bs.acr.2018.02.006

30. Jan, Z., Ahmed, W. S., Biswas, K. H., & Jithesh, P. V. (2023). Identification of a potential DNA methyltransferase (DNMT) inhibitor. Journal of biomolecular structure & dynamics, 1–15. Advance online publication. 10.1080/07391102.2023.2233637

31. Marzochi, L. L., Cuzziol, C. I., Nascimento Filho, C. H. V. D., Dos Santos, J. A., Castanhole-Nunes, M. M. U., Pavarino, É. C., Guerra, E. N. S., & Goloni-Bertollo, E. M. (2023). Use of histone methyltransferase inhibitors in cancer treatment: A systematic review. European journal of pharmacology, 944, 175590. 10.1016/j.ejphar.2023.175590

32. Baby, S., Gurukkala Valapil, D., & Shankaraiah, N. (2021). Unravelling KDM4 histone demethylase inhibitors for cancer therapy. Drug discovery today, 26(8), 1841–1856. 10.1016/j.drudis.2021.05.015

33. Gajjela, B. K., & Zhou, M. M. (2023). Bromodomain inhibitors and therapeutic applications. Current opinion in chemical biology, 75, 102323. 10.1016/j.cbpa.2023.102323

34. Yunfeng Qi, Dadong Wang, Daying Wang, Taicheng Jin, Liping Yang, Hui Wu, Yaoyao Li, Jing Zhao, Fengping Du, Mingxia Song, and Renjun Wang HEDD: the human epigenetic drug database Database, 2016: pii: baw159.

35. Shailesh, H., Zakaria, Z. Z., Baiocchi, R., & Sif, S. (2018). Protein arginine methyltransferase 5 (PRMT5) dysregulation in cancer. Oncotarget, 9(94), 36705–36718.

36. Xiao, W., Chen, X., Liu, L., Shu, Y., Zhang, M., & Zhong, Y. (2019). Role of protein arginine methyltransferase 5 in human cancers. Biomedicine and Pharmacotherapy, 114:108790.

37. Tremblay-Lemay, R., Rastgoo, N., Pourabdollah, M., & Chang, H. (2018). EZH2 as a therapeutic target for multiple myeloma and other haematological malignancies. Biomarker Research, 6:34.

38. Gan, L., Yang, Y., Li, Q., Feng, Y., Liu, T., & Guo, W. (2018). Epigenetic regulation of cancer progression by EZH2: From biological insights to therapeutic potential. Biomarker Research, 6:10.

39. Eich, M. L., Athar, M., Ferguson, J. E., 3rd, & Varambally, S. (2020). EZH2-Targeted Therapies in Cancer: Hype or a Reality. Cancer research, 80(24), 5449–5458. 10.1158/0008-5472.CAN-20-2147

40. Rudenko, A. Y., Mariasina, S. S., Sergiev, P. V., & Polshakov, V. I. (2022). Analogs of *S*-Adenosyl-*L*-Methionine in Studies of Methyltransferases. Molecular biology, 56(2), 229–250. 10.1134/S002689332202011X

41. Song, X., Gao, T., Wang, N., Feng, Q., You, X., Ye, T., Lei, Q., Zhu, Y., Xiong, M., Xia, Y., Yang, F., Shi, Y., Wei, Y., Zhang, L., & Yu, L. (2016). Selective inhibition of EZH2 by ZLD1039 blocks H3K27 methylation and leads to potent anti-tumor activity in breast cancer. Scientific reports, 6, 20864. 10.1038/srep20864

42. Wang, Q., Xu, J., Li, Y., Huang, J., Jiang, Z., Wang, Y., Liu, L., Leung, E. L. H., & Yao, X. (2018). Identification of a Novel Protein Arginine Methyltransferase 5 Inhibitor in Non-small Cell Lung Cancer by Structure-Based Virtual Screening. Frontiers in pharmacology, 9, 173. 10.3389/fphar.2018.00173

43. Yang, L., Ma, D. W., Cao, Y. P., Li, D. Z., Zhou, X., Feng, J. F., & Bao, J. (2021). PRMT5 functionally associates with EZH2 to promote colorectal cancer progression through epigenetically repressing CDKN2B expression. Theranostics, 11(8), 3742–3759. 10.7150/thno.53023

44. Kedhari Sundaram, M., Haque, S., Somvanshi, P., Bhardwaj, T., & Hussain, A. (2020). Epigallocatechin gallate inhibits HeLa cells by modulation of epigenetics and signaling pathways. 3 Biotech, 10(11), 484. 10.1007/s13205-020-02473-1

45. Balamurugan, K., & Pisabarro, M. T. (2021). Stabilizing Role of Water Solvation on Anion-π Interactions in Proteins. ACS omega, 6(39), 25350–25360. 10.1021/acsomega.1c03264

46. Chen, Y., Shao, X., Zhao, X., Ji, Y., Liu, X., Li, P., Zhang, M., & Wang, Q. (2021). Targeting protein arginine methyltransferase 5 in cancers: Roles, inhibitors and mechanisms. In Biomedicine & Pharmacotherapy (Vol. 144, p. 112252). Elsevier BV. 10.1016/j.biopha.2021.112252

47. Rasheed, Z., Rasheed, N., & Al-Shaya, O. (2018). Epigallocatechin-3-O-gallate modulates global microRNA expression in interleukin-1β-stimulated human osteoarthritis chondrocytes: potential role of EGCG on negative co-regulation of microRNA-140-3p and ADAMTS5. European Journal of Nutrition, 57(3), 917–928.

48. Shankar, S., Marsh, L., & Srivastava, R. K. (2013). EGCG inhibits growth of human pancreatic tumors orthotopically implanted in Balb C nude mice through modulation of FKHRL1/FOXO3a and neuropilin. Molecular and Cellular Biochemistry, 372(1–2), 83–94.

49. Balasubramanian Sivaprakasam, Gautam Adhikary, Richard L. Eckert, The Bmi-1 polycomb protein antagonizes the (−)-epigallocatechin-3-gallate-dependent suppression of skin cancer cell survival Carcinogenesis, Volume 31, Issue 3, March 2010, Pages 496–503,

50. Li, Y., Yang, Y., Liu, X., Long, Y., & Zheng, Y. (2019). PRMT5 Promotes Human Lung Cancer Cell Apoptosis via Akt/Gsk3β Signaling Induced by Resveratrol. Cell transplantation, 28(12), 1664–1673. 10.1177/0963689719885083

51. Xia, T., Liu, M., Zhao, Q., Ouyang, J., Xu, P., & Chen, B. (2021). PRMT5 regulates cell pyroptosis by silencing CASP1 in multiple myeloma. Cell death & disease, 12(10), 851. 10.1038/s41419-021-04125-5

52. Vlummens, P., Verhulst, S., De Veirman, K., Maes, A., Menu, E., Moreaux, J., De Boussac, H., Robert, N., De Bruyne, E., Hose, D., Offner, F., Vanderkerken, K. & Maes, K. (2022). Inhibition of the Protein Arginine Methyltransferase PRMT5 in High-Risk Multiple Myeloma as a Novel Treatment Approach. Frontiers in cell and developmental biology, 10, 879057. 10.3389/fcell.2022.879057

53. Kong, J., Wang, Z., Zhang, Y., Wang, T., & Ling, R. (2023). Protein Arginine Methyltransferases 5 (PRMT5) affect Multiple Stages of Autophagy and Modulate Autophagy-related Genes in Controlling Breast Cancer Tumorigenesis. Current cancer drug targets, 23(3), 242–250. 10.2174/1568009622666220922093059

54. Brobbey, C., Yin, S., Liu, L., Ball, L. E., Howe, P. H., Delaney, J. R., & Gan, W. (2023). Autophagy dictates sensitivity to PRMT5 inhibitor in breast cancer. Scientific Reports, 13(1), 1–13. 10.1038/s41598-023-37706-9

55. Liu, J. H., Cao, Y. M., Rong, Z. P., Ding, J., & Pan, X. (2021). Trichostatin A Induces Autophagy in Cervical Cancer Cells by Regulating the PRMT5-STC1-TRPV6-JNK Pathway. Pharmacology, 106(1-2), 60–69. 10.1159/000507937

56. Gupta, S., Hussain, T., & Mukhtar, H. (2003). Molecular pathway for (-)-epigallocatechin-3-gallate-induced cell cycle arrest and apoptosis of human prostate carcinoma cells. Archives of biochemistry and biophysics, 410(1), 177–185. 10.1016/s0003-9861(02)00668-9

57. Zhou, D. H., Wang, X., Yang, M., Shi, X., Huang, W., & Feng, Q. (2013). Combination of low concentration of (-)-epigallocatechin gallate (EGCG) and curcumin strongly suppresses the growth of non-small cell lung cancer in vitro and in vivo through causing cell cycle arrest. International journal of molecular sciences, 14(6), 12023–12036. 10.3390/ijms140612023

58. Kumar, N. B., Pow-Sang, J., Spiess, P. E., Park, J., Salup, R., Williams, C. R., Parnes, H., & Schell, M. J. (2016). Randomized, placebo-controlled trial evaluating the safety of one-year administration of green tea catechins. Oncotarget, 7(43), 70794–70802. 10.18632/oncotarget.12222

59. Gee, J. R., Saltzstein, D. R., Kim, K., Kolesar, J., Huang, W., Havighurst, T. C., Wollmer, B. W., Stublaski, J., Downs, T., Mukhtar, H., House, M. G., Parnes, H. L., & Bailey, H. H. (2017). A Phase II Randomized, Double-blind, Presurgical Trial of Polyphenon E in Bladder Cancer Patients to Evaluate Pharmacodynamics and Bladder Tissue Biomarkers. Cancer prevention research (Philadelphia, Pa.), 10(5), 298–307. 10.1158/1940-6207.CAPR-16-0167

60. Choudhari, A. S., Mandave, P. C., Deshpande, M., Ranjekar, P., & Prakash, O. (2020). Phytochemicals in Cancer Treatment: From Preclinical Studies to Clinical Practice. Frontiers in pharmacology, 10, 1614. 10.3389/fphar.2019.01614

61. Cerami, E., Gao, J., Dogrusoz, U., Gross, B. E., Sumer, S. O., Aksoy, B. A., Jacobsen, A., Byrne, C. J., Heuer, M. L., Larsson, E., Antipin, Y., Reva, B., Goldberg, A. P., Sander, C., & Schultz, N. (2012). The cBio cancer genomics portal: an open platform for exploring multidimensional cancer genomics data. Cancer discovery, 2(5), 401–404. 10.1158/2159-8290.CD-12-0095 (https://www.cbioportal.org/)

62. Tsai, C. Y., Chen, C. Y., Chiou, Y. H., Shyu, H. W., Lin, K. H., Chou, M. C., Huang, M. H., & Wang, Y. F. (2017). Epigallocatechin-3-Gallate Suppresses Human Herpesvirus 8 Replication and Induces ROS Leading to Apoptosis and Autophagy in Primary Effusion Lymphoma Cells. International journal of molecular sciences, 19(1), 16. 10.3390/ijms19010016

63. Zhang, S., Cao, M., & Fang, F. (2020). The Role of Epigallocatechin-3-Gallate in Autophagy and Endoplasmic Reticulum Stress (ERS)-Induced Apoptosis of Human Diseases. Medical science monitor : international medical journal of experimental and clinical research, 26, e924558. 10.12659/MSM.924558

